# Atopic Dermatitis and Psoriasis Differ in Lesional DEG Reference Instability and Non-Lesional Spectrum Displacement: Multi-Cohort Geometric Evidence of Individual Homeostatic Boundary Escape

**DOI:** 10.64898/2026.03.11.711058

**Authors:** Basma Shabana

**Author notes:** <u>Email:</u>.

## Abstract

Atopic dermatitis (AD) transcriptomic studies have produced notoriously inconsistent differential gene expression (DEG) lists across cohorts, and the biological heterogeneity of non-lesional AD skin remains unresolved at the individual level --limitations that group-averaged DEG analysis is structurally incapable of addressing. We applied a geometric transcriptomic framework to 537 skin RNA-sequencing samples from four independent international cohorts, positioning each sample within a 20-dimensional disease-informative PCA space relative to a unified healthy reference cloud validated by multi-cohort batch integration (ComBat; silhouette width 0.076 across three countries).

Applying a bootstrap Jaccard framework to lesional-versus-healthy comparisons, AD lesional DEG signatures were significantly more reference-sensitive than psoriasis (PSO; Jaccard 0.637 vs 0.737; non-overlapping 95% CIs), with per-gene Spearman correlation between absolute effect size and cross-reference reproducibility (rho=0.658, p<2.2x10-16) establishing that AD’s instability arises from its structural dependence on smaller-effect-size transcriptomic signals -- a geometric property of AD biology, not a methodological failure of prior studies.

In non-lesional skin, spectrum scores -- projections of each sample’s displacement from the healthy centroid onto the disease axis -- revealed that AD non-lesional (AD_NL) samples had traversed 22.6% (95% CI 15.2-29.6%) of the healthy-to-lesional axis versus 12.9% (95% CI 8.4-18.2%) for PSO non-lesional (PSO_NL; Wilcoxon p=0.012), with non-lesional skin in each disease displaced toward its own lesional pole at near-identical angles (within-disease delta-angle 1.4 degrees) but AD displaced further. Against a homeostatic boundary defined as the 95th-percentile Euclidean distance of healthy controls from their centroid, 17.2% of AD_NL samples (16/93) individually exceeded this threshold versus 2.0% of PSO_NL (1/49; OR=9.87, p=0.007), replicating across all three cohorts providing AD_NL data. Among boundary-crossing AD_NL samples, two directional patterns emerged: high-positive displacement (n=6) characterised by inflammatory pre-activation (IL-6/JAK-STAT3, IFN-alpha, TNF-alpha/NF-kB) and low/negative displacement (n=10) characterised by broad metabolic suppression and a third transcriptomic axis orthogonal to both canonical disease trajectories -- not attributable to cellular infiltration differences and undetectable by group-level analysis.

These findings reframe AD’s notorious transcriptomic inconsistency as a predictable consequence of effect-size architecture, establish that a reproducible subset of AD patients harbours individually measurable transcriptomic boundary escape before clinical lesion onset, and identify a biologically uncharacterised non-lesional subgroup that warrants cell-type-resolved investigation as a potential early intervention target.

## Introduction

Atopic dermatitis (AD) and psoriasis (PSO) represent the two most extensively studied chronic inflammatory skin diseases, yet fundamental questions about their subclinical biology remain unresolved. Early comparative studies highlighted stark pathogenic differences: AD is characterized by Th2-skewed immunity with prominent barrier defects, whereas PSO is driven by IL-17/Th17 pathways with epidermal hyperplasia [1,2]. These distinctions extend to non-lesional skin, where clinical observations suggest AD exhibits dryness, pruritus, and barrier compromise even in unaffected areas, while PSO non-lesional skin appears largely normal [3,4].

Transcriptomic profiling has progressively refined these insights. Initial bulk RNA-sequencing analyses revealed that non-lesional AD skin displays mild but detectable pro-inflammatory and epidermal changes—such as downregulation of filaggrin (FLG) and late cornified envelope genes, alongside Th2/Th17/Th22 skewing—that intensify in lesional sites [5,6]. In contrast, non-lesional PSO skin shows minimal deviation from healthy controls, with only subtle IL-17-related signals [7,8]. Larger-scale studies confirmed greater molecular heterogeneity in AD compared to PSO, including variable Th1/Th2/Th17/Th22 activation across subtypes, ages, and ethnicities, contributing to inconsistent DEG lists across independent cohorts [5,9]. Recent unbiased multi-cohort analyses have further underscored this disparity: non-lesional AD exhibits weaker but lesional-like immune and barrier dysregulation, while non-lesional PSO profiles converge closely with healthy skin [10,11].

Despite these advances, standard differential gene expression (DEG) analysis—the dominant framework in skin transcriptomics—remains constrained by two key limitations. First, it is inherently sensitive to the composition of the healthy reference cohort, as healthy skin varies across populations in immune tone, microbiome exposure, and epidermal barrier expression; this reference dependency can yield poorly overlapping DEG lists even when studies converge on shared mechanisms [12,13]. Second, group-averaged signatures obscure individual-level variation, precluding the characterization of qualitatively distinct non-lesional abnormalities that may reflect mechanistic heterogeneity or predict therapeutic response [14,15].

A deeper conceptual issue underlies the reference sensitivity problem. Differential expression analysis implicitly treats disease as a binary contrast: genes are either significantly displaced from the healthy reference or they are not. When the molecular signal is large — as in psoriatic lesional skin, where IL-17-driven remodelling displaces thousands of genes by substantial margins — this binary framing is adequate, because the magnitude of displacement greatly exceeds any reasonable variation in where the healthy reference is positioned. However, when molecular changes are distributed across many genes with individually modest effects, as may characterize diseases involving gradual subclinical remodelling rather than switch-like activation, the detected signature becomes sensitive to precisely where the healthy reference is placed. In this regime, the reference is not merely a technical convenience but a structural determinant of the result: genes whose disease-associated shift is small relative to the within-healthy distribution will cross significance thresholds differently depending on which individuals constitute the reference cohort, even after batch correction removes technical variation. Reconceptualising disease biology as a continuous spectrum between healthy and lesional poles — rather than a binary deviation from a fixed zero — reframes this problem productively. Within such a framework, signal proximity to the healthy pole predicts reference sensitivity: genes far from healthy are reproducibly detected regardless of reference composition; genes close to healthy are marginal and therefore reference-dependent. This geometric interpretation explains why reference sensitivity should be expected to correlate with effect size, and why diseases with more distributed, lower-amplitude molecular signatures should exhibit greater inter-cohort DEG variability than diseases with large, coherent effector responses.

To overcome these limitations, we developed a geometric analytical framework that positions each individual sample within a multi-dimensional transcriptomic space relative to a unified healthy reference cloud. By defining a homeostatic boundary as the 95th-percentile Euclidean distance from the healthy centroid in a disease-informed principal component space, we identify samples that have departed from normal homeostasis. Critically, sensitivity analyses demonstrate that a coordinate system derived solely from healthy variance fails to resolve subtle pre-disease signals in non-lesional AD — capturing florid lesional disease in both conditions while leaving subclinical dysregulation below the threshold of detection. This empirical limitation motivates the inclusion of disease-axis variance in the principal component space: gene programs active in early AD (e.g., CXCL8, FLG2, CCL20, RORC) show negligible variation within healthy skin and are therefore underweighted in a healthy-only decomposition, rendering the resulting axes insensitive to the very signals this study aims to characterize [5,10].

Applied to an integrated dataset of 537 skin transcriptomes from four independent international cohorts, this framework quantifies individual-level transcriptomic displacement and directional heterogeneity in non-lesional skin. We hypothesized that AD non-lesional samples would exhibit greater and more variable departure from healthy homeostasis than PSO non-lesional samples, reflecting AD’s proposed continuum of subclinical priming versus PSO’s more switch-like transcriptomic transition. The analysis reveals disease-specific patterns of boundary escape and spectrum positioning, providing multi-cohort, individual-level geometric evidence of transcriptomic pre-lesional states in AD, mechanistic insight into reference sensitivity, and hypothesis-generating directional patterns for future endotype and therapeutic stratification studies.

## Methods

### Datasets and Sample Selection

We integrated raw RNA-sequencing count data from four publicly available Gene Expression Omnibus (GEO) datasets spanning diverse geographic origins and clinical phenotypes: GSE121212 (United States; AD, PSO, and healthy controls), GSE193309 (Denmark; GENAD cohort, AD and healthy controls), GSE306437 (Ireland; AD and PSO), and GSE54456 (United States; PSO lesional and healthy controls). These datasets provide featureCounts-quantified reads aligned to GRCh38/NCBI.

To prevent pseudoreplication in longitudinal data, only baseline visit samples (visit identifier ‘01’) were retained from GSE193309 (reducing 339 to 148 samples). From GSE306437, only week-1 (V1) samples were included; week-12 post-dupilumab samples (V12) were excluded (reducing 118 to 75 samples). After systematic quality control and outlier removal, the final analytical dataset comprised 537 samples across five groups: healthy controls (n=166), AD non-lesional (AD_NL; n=93), AD lesional (AD_LS; n=90), PSO non-lesional (PSO_NL; n=49), and PSO lesional (PSO_LS; n=139) (Table 1).

**Table 1:** Sample composition of the final analytical dataset after quality control and outlier removal. Rows = four GEO cohorts (GSE121212, GSE193309, GSE306437, GSE54456); columns = five biological groups (Healthy, AD_NL, AD_LS, PSO_NL, PSO_LS). Sample counts: Healthy = 166, AD_NL = 93, AD_LS = 90, PSO_NL = 49, PSO_LS = 139; total n = 537. GSM5788875 (|PC1 z-score| > 3) was excluded from PCA training but projected into the coordinate system for downstream analyses; all geometric analyses including boundary crossing and spectrum scores are performed on n = 537.

| Dataset | Healthy | AD_NL | AD_LS | PSO_NL | PSO_LS |
| --- | --- | --- | --- | --- | --- |
| **GSE121212** | 37 | 27 | 26 | 27 | 27 |
| **GSE193309** | 48 | 51 | 48 | 0 | 0 |
| **GSE306437** | 0 | 15 | 16 | 22 | 22 |
| **GSE54456** | 81 | 0 | 0 | 0 | 90 |
| **TOTAL** | 166 | 93 | 90 | 49 | 139 |

### Metadata Harmonisation and Group Classification

Metadata were extracted from GEO SOFT files using a custom parser and harmonized into five unified group labels: Healthy, AD_NL, AD_LS, PSO_NL, and PSO_LS. AD_Chronic samples in GSE121212 (n=6) were reclassified as AD_LS, consistent with clinical definitions of chronic lesional disease. For GSE306437, GTG library codes were mapped to GSM identifiers via a parsed library-name table.

### Gene Identifier Harmonisation and Quality Filtering

Entrez gene identifiers were converted to HGNC symbols using the org.Hs.eg.db Bioconductor annotation package (v3.17). Duplicate mappings were resolved by retaining the first symbol; genes mapping to the same symbol were collapsed by summing raw counts. Low-abundance genes were filtered per dataset (minimum count of 10 in ≥20% of samples and in at least three samples). The four filtered gene sets were intersected, yielding 15,925 genes measurable across all datasets.

### Normalisation and Batch Correction

Variance-stabilising transformation (VST) was applied independently to each dataset using DESeq2 (v1.42) with blind=TRUE and an intercept-only design [16]. This per-dataset VST ensures dispersion estimates reflect within-dataset variation only. The four VST matrices were merged and batch-corrected using ComBat from the sva package (v3.50), applying empirical Bayes adjustment to the VST-transformed data with batch defined as study of originand a protection matrix defined as model.matrix(∼ group) incorporating the five-level group factor (Healthy, AD_NL, AD_LS, PSO_NL, PSO_LS) [17]. For datasets with unrepresented group levels (GSE54456 contributes no PSO_NL; GSE306437 contributes no healthy controls), ComBat’s empirical Bayes estimation uses the available group terms, with the global pooled prior stabilising estimates for sparse cells. This approach preserves biological signals while mitigating technical confounding.

### Reference Sensitivity Analysis

Healthy controls were clustered in Seurat (FindNeighbors + FindClusters, resolution=0.5). Silhouette widths were computed on scaled PCA distances using the cluster package [19]. DEG analysis between each disease’s lesional samples and each healthy sub-cluster was performed using DESeq2 with study as covariate and |log2FC| >0.5 [16]. Reference sensitivity was quantified as the Jaccard similarity index between cluster-specific DEG lists. Spearman correlation between per-gene Jaccard reproducibility and absolute log2FC was computed across all tested genes.

### PCA Coordinate System and Feature Space

Following PCA outlier exclusion from training (samples with |PC1 z-score| >3; n=1, GSM5788875), the cleaned 537-sample matrix was used to fit the PCA coordinate system in Seurat (v5.0) [18]. The 3,000 most variable features were selected from the full mixed dataset, data were scaled to unit variance per gene, and 20 principal components were computed (RunPCA, npcs=20). The resulting PCA rotation was then used to project all 537 samples (including GSM5788875) into the 20-PC coordinate system (pca_coords) for all downstream geometric analyses.

Feature selection was validated by comparing overlap with the top 3,000 variable genes from healthy controls alone (33.8% overlap). The 888 genes unique to the mixed set — including CXCL8, FLG2, CCL20, RORC, DSC2, and IL34 — show negligible variance within healthy skin (range 0.13–0.24) but are highly variable across the mixed dataset (variance 1.13–4.22), explaining why they are underweighted in a healthy-only decomposition(Supplementary Table S1).

A formal sensitivity analysis using a healthy-only PCA confirmed that this coordinate system is empirically necessary for detecting subtle pre-disease activation: boundary-crossing was non-significant in AD_NL under the healthy-only space (OR=2.15, p=0.659), confirming that a healthy-derived coordinate system lacks power to detect subclinical dysregulation, while florid lesional disease was robustly identified in both conditions (AD_LS OR=6.65, p<0.001; PSO_LS OR=70.47, p<0.001)(Supplementary Figure S1).

The mixed-feature space is therefore biologically motivated and empirically supported.

### Homeostatic Boundary and Distance Analysis

The homeostatic boundary was defined as the 95th-percentile Euclidean distance of healthy controls from their centroid in 20-dimensional PCA space (radius=53.74 PCA units). Group centroids were computed as column means within each group. Normalised distances were calculated as each sample’s distance from the healthy centroid divided by this radius. Boundary-crossing significance was assessed using three Fisher exact tests (AD_NL vs PSO_NL, AD_NL vs Healthy, PSO_NL vs Healthy) and a permutation test (n=10,000 label permutations of AD_NL and PSO_NL).

### Geometric Trajectory Analysis

Disease axis vectors were defined as the vector from the healthy centroid to each disease’s lesional centroid. Spectrum scores were computed as the scalar projection of each sample’s displacement from the healthy centroid onto the disease axis unit vector, expressed as a percentage of the total healthy-to-lesional axis length. Values exceeding 100% or falling below 0% indicate projection beyond the lesional centroid or in the opposing direction, respectively, and are retained as informative extremes. Cosine similarity between healthy-to-non-lesional and healthy-to-lesional vectors was computed using the dot product formula. Bootstrap 95% confidence intervals (n=1,000 resamples) were generated for group-level spectrum scores.

### Directional Pattern Analysis and Formal Bimodality Testing

Among boundary-crossing AD_NL samples (n=16), directional patterns were characterised by spectrum score. Samples with spectrum score >30% were classified as high-positive displacement (toward lesional); those ≤30% as low/negative displacement (lateral). The 30% threshold corresponds to the first quartile of V1 spectrum scores in GSE306437 and was validated as stable across 20–50% thresholds; it should be interpreted as an exploratory, sensitivity-validated partition rather than a definitive biological boundary.

Formal bimodality was assessed using Hartigan’s dip test (diptest package, B=10,000 simulations) [20] and Gaussian mixture modelling (mclust package, G=1–4) [21]. Results are reported as observational and hypothesis-generating given the small sample size.

### Pathway Analysis

Pathway activity was scored for all 537 samples using GSVA (v1.50) with 50 MSigDB Hallmark gene sets (v2023.2) and kcdf=’Gaussian’ (appropriate for continuous VST data) [22,23]. Differential pathway activity between groups was tested using Wilcoxon rank-sum tests with Benjamini-Hochberg correction across 50 pathways.

### Cell Type Deconvolution

Immune cell proportions were estimated using the EPIC algorithm (immunedeconv R package v2.1) from batch-corrected expression data [24]. Three samples showed incomplete optimisation convergence (all lesional; none contributed to AD_NL boundary analysis). Two pre-specified Wilcoxon comparisons were performed: (i) all boundary-crossing AD_NL samples (n=16) versus within-boundary AD_NL samples (n=77), to test whether boundary escape is associated with altered cellular composition overall; and (ii) a three-way comparison of low/negative-displacement (n=10), high-positive-displacement (n=6), and within-boundary AD_NL (n=77), to test whether the two directional subgroups differ in cellular composition from each other or from within-boundary samples.

### Robustness Analyses

Five robustness tests were performed: (i) PC sensitivity at 10, 15, and 20 PCs; (ii) Mahalanobis distance boundary replacing Euclidean radius; (iii) permutation testing (n=10,000); (iv) directional-pattern threshold sensitivity at 20%, 30%, 40%, and 50%; (v) GSVA pathway robustness after removing the most extreme sample (GSM5788875, spectrum score=−126.3%, reflecting projection in the direction opposite to the disease axis at greater magnitude than the axis length itself).

### Statistical Analysis

Fisher exact tests used mid-P estimates. Bootstrap confidence intervals are 95th-percentile intervals from n=1,000 resamples. Permutation p-values represent the fraction of 9,993 valid permutations with OR ≥ observed OR. All analyses were performed in R (v4.3).

## Results

### Healthy Skin from Three International Cohorts Converges into a Single Homogeneous Reference Space

Following ComBat batch correction applied to the 537 samples from four independent studies, the 166 healthy control samples formed a coherent, homogeneous cluster whose dispersion showed no meaningful structuring by cohort of origin. Unsupervised clustering of healthy controls identified two to three small sub-clusters with no substantial structure (average silhouette width = 0.076; values below 0.1 indicate absence of meaningful clustering), and visual inspection confirmed that residual structure primarily reflected study of origin rather than biological subpopulations [18,19]. Prior to batch correction, the four datasets occupied distinct regions of PCA space; after correction, samples segregated by disease group, while healthy controls from the United States, Denmark, and Ireland interdigitated freely (Figure 1). This convergence establishes a single, valid centroid and radius for the healthy reference cloud, enabling interpretable inter-sample distances.

**Figure 1:**
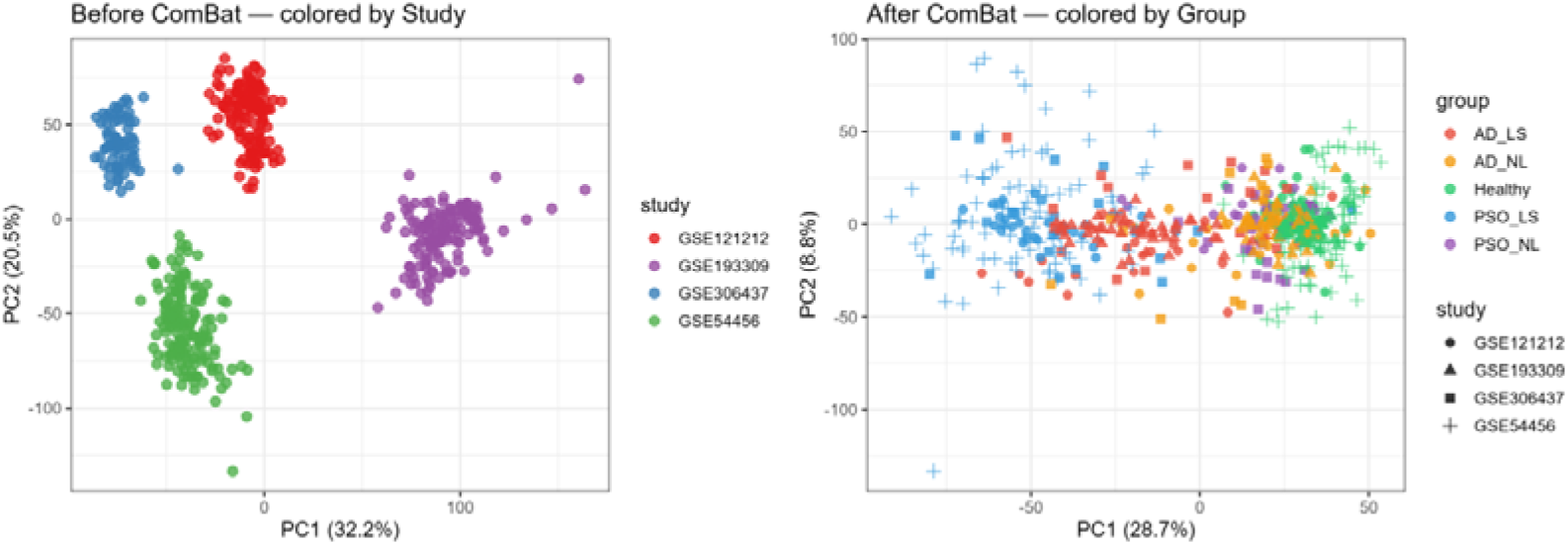
PCA plots of 537 skin transcriptomes before and after ComBat batch correction. Left: Prior to correction, samples segregate completely by study of origin (PC1 32.2%, PC2 20.5%), with each of the four GEO datasets (GSE121212, GSE193309, GSE306437, GSE54456) occupying a distinct, non-overlapping region of PCA space. Right: Following empirical Bayes ComBat correction with a group-protected design matrix (model.matrix(∼ group), five-level factor), samples reorganize by biological group. Healthy controls from the four cohorts (circles = GSE121212; triangles = GSE193309; squares = GSE306437; crosses = GSE54456) interdigitate freely, forming a single coherent reference cloud. Disease groups (AD_NL, AD_LS, PSO_NL, PSO_LS) are distinguishable by biological group, confirming that batch correction preserved disease signal while removing technical confounding. PC1 and PC2 explain 28.7% and 8.8% of post-correction variance, respectively. One outlier (GSM5788875, |PC1 z-score| > 3) was excluded prior to all geometric analyses; final analytical n = 537.

### AD Differential Expression Signatures Are Significantly More Reference-Sensitive Than PSO

To quantify the impact of healthy reference composition on disease DEG lists, we implemented a bootstrap Jaccard framework. Healthy controls were divided into sub-clusters by unsupervised analysis, and DEG analysis was repeated independently using each cluster as reference. The Jaccard index was 0.637 for AD (range 0.561–0.728; bootstrap 95% CI 0.612–0.661) and 0.737 for PSO (range 0.680–0.821; 95% CI 0.718–0.756; Figure 2A). Non-overlapping bootstrap 95% CIs indicate a statistically significant group difference, confirming that AD signatures are significantly more reference-sensitive: approximately 36% of the AD DEG list changes with reference composition, compared with 26% for PSO.

**Figure 2A:**
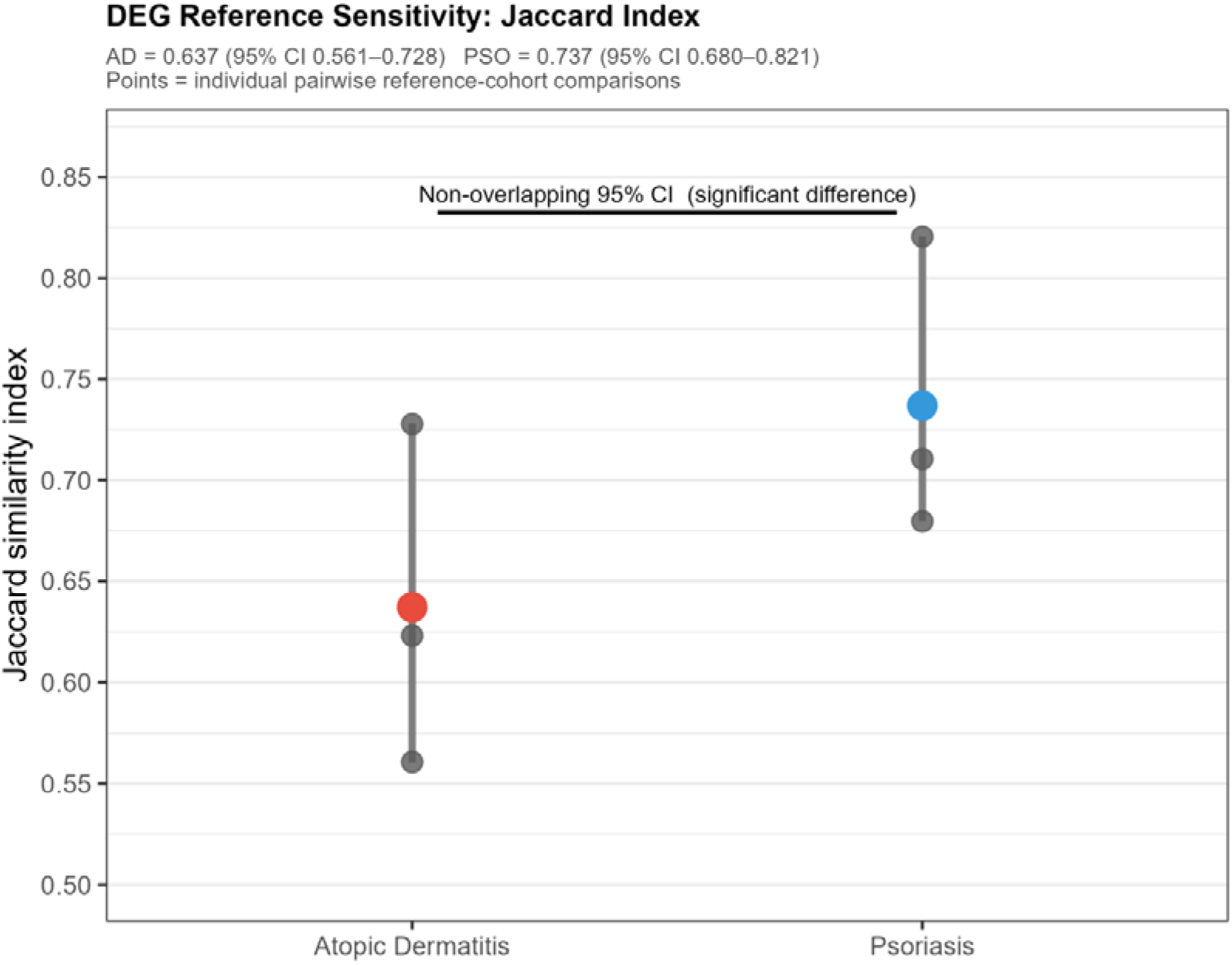
DEG reference sensitivity quantified by the Jaccard similarity index for atopic dermatitis (AD, red) and psoriasis (PSO, blue). Each grey point represents one pairwise Jaccard comparison between DEG lists generated using different healthy sub-cluster references (three comparisons per disease). The coloured point is the group mean; vertical grey segment spans the observed range. The horizontal bracket confirms non-overlapping bootstrap 95% confidence intervals (B = 1,000 resamples, set.seed(42)), indicating a statistically significant difference between diseases. Mean Jaccard: AD = 0.637 (observed range 0.561–0.728; bootstrap 95% CI 0.612–0.661); PSO = 0.737 (observed range 0.680–0.821; bootstrap 95% CI 0.718–0.756). A higher Jaccard index reflects greater stability of the DEG list across reference choices. Approximately 36% of AD DEG genes change with reference composition versus 26% for PSO, indicating that AD transcriptomic signatures are significantly more dependent on healthy reference composition than PSO signatures.

Reference-invariant (core) AD genes (n=1,583) were enriched for epidermal barrier function (FLG, LOR, SPRR1A/B, KRT1/10, TGM1, CLDN1), whereas reference-sensitive genes (n=1,565) were enriched for GTPase signalling and epithelial-mesenchymal transition. For PSO, the core set (n=3,434) comprised barrier and keratinocyte proliferation genes (KRT6A/16/17, S100A7/8/9, DEFB4A, IL36G, CXCL1/8), with sensitive genes (n=2,025) enriched for mitochondrial metabolism (Table 2).

**Table 2:**
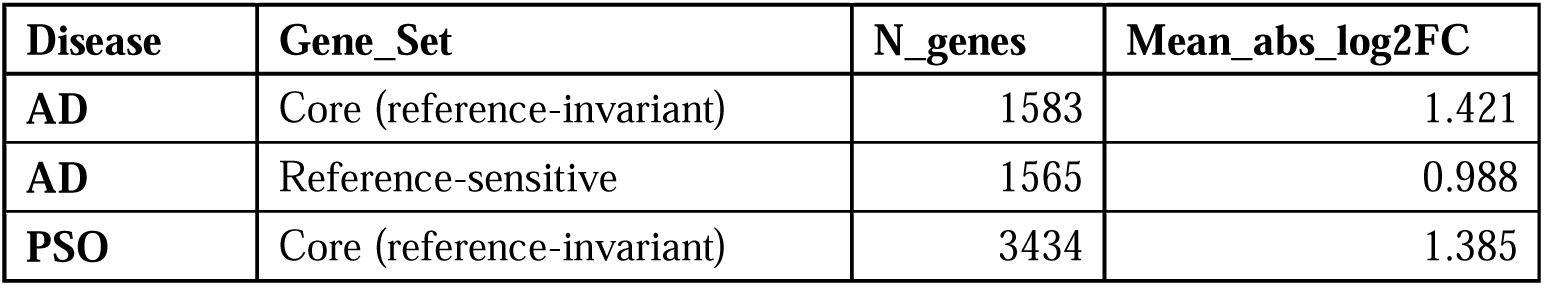
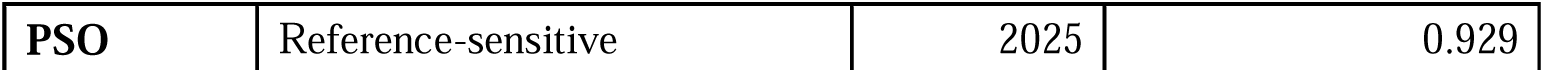
Mean absolute log□ fold-change for reference-invariant (core) and reference-sensitive gene sets in AD and PSO. Core AD genes (n = 1,583): mean |log□FC| = 1.421; reference-sensitive AD genes (n = 1,565): mean |log□FC| = 0.988. Core PSO genes (n = 3,434): mean |log□FC| = 1.385; reference-sensitive PSO genes (n = 2,025): mean |log□FC| = 0.929. In both diseases, core (reference-invariant) genes show approximately 0.4–0.5 higher mean effect size than reference-sensitive genes, directly supporting the effect-size mechanism for reference sensitivity documented in Figure 2B.

| Disease | Gene_Set | N_genes | Mean_abs_log2FC |
| --- | --- | --- | --- |
| AD | Core (reference-invariant) | 1583 | 1.421 |
| AD | Reference-sensitive | 1565 | 0.988 |
| PSO | Core (reference-invariant) | 3434 | 1.385 |
| PSO | Reference-sensitive | 2025 | 0.929 |

Effect size emerged as the primary driver of reference sensitivity: core AD genes exhibited a mean |log2FC| of 0.965 versus 0.611 for sensitive genes (PSO: 1.053 vs. 0.600). Across all tested genes, Spearman correlation between per-gene effect size and cross-cluster reproducibility was rho=0.687 (p<2.2×10□¹□; Figure 2B). Thus, AD reference instability arises directly from reliance on smaller-effect-size transcriptomic signals, consistent with AD’s documented molecular heterogeneity [5,9,10].

**Figure 2B:**
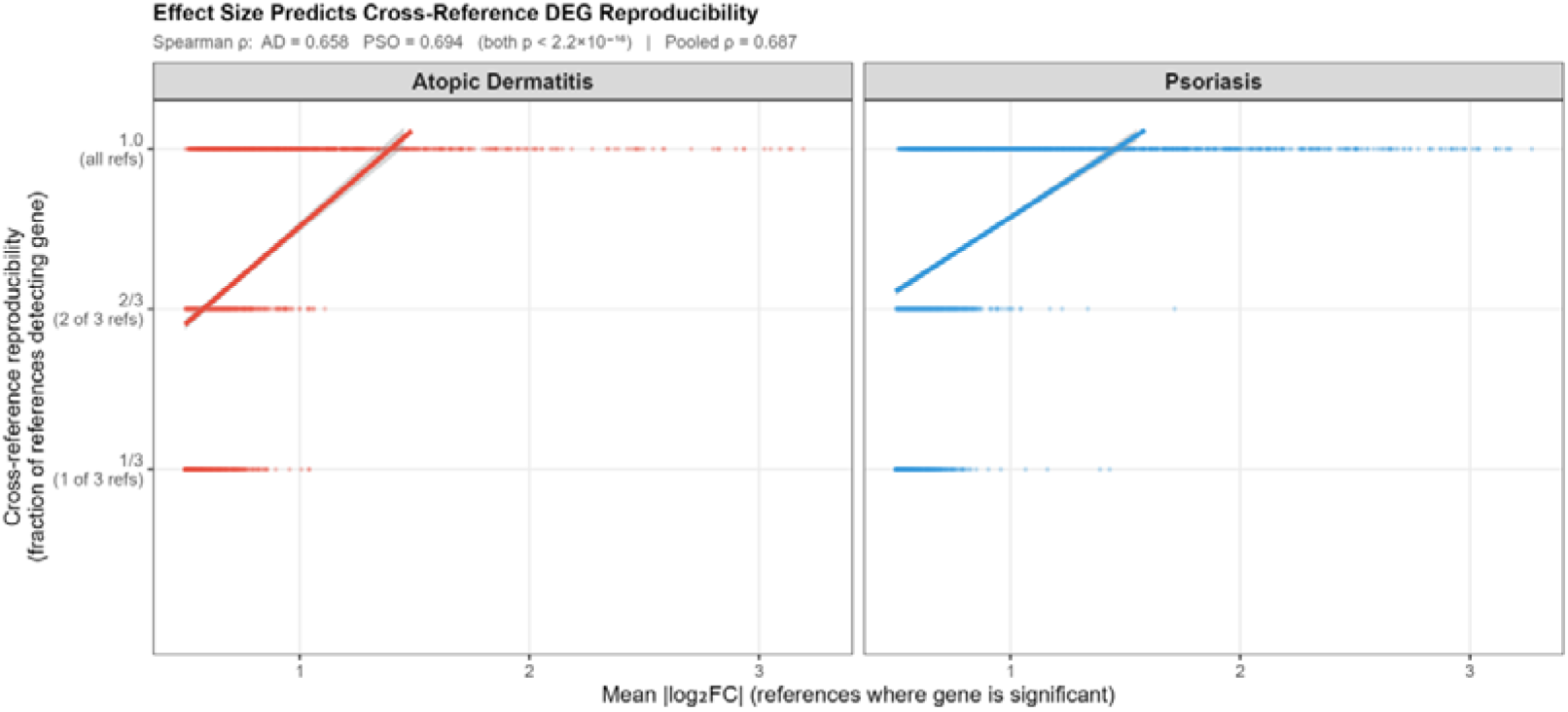
Scatter plots of per-gene mean absolute log□ fold-change (x-axis) versus cross-reference reproducibility (y-axis: fraction of reference cohorts in which the gene reached DEG significance at padj < 0.05, |log□FC| > 0.5; values: 1/3, 2/3, or 1.0 for three reference comparisons) for atopic dermatitis (AD, red, left panel) and psoriasis (PSO, blue, right panel). Analysis is restricted to genes significant in at least one reference cohort; genes never detected across any reference carry no information about reference sensitivity and are excluded. Each point is one gene; the regression line with 95% confidence band is fitted by ordinary least squares. The x-axis is capped at the 99th percentile of |log□FC| to prevent extreme outliers compressing the main cloud. In both diseases, higher effect size is associated with more consistent detection across reference cohorts, reflected in a positive regression slope and positive Spearman ρ: AD = 0.658; PSO = 0.694 (both p < 2.2×10□¹□); pooled across both diseases ρ = 0.687 (p < 2.2×10□¹□). The three discrete y-axis levels reflect the ordinal nature of the reproducibility variable with three reference comparisons. This figure demonstrates that AD reference instability arises primarily from reliance on smaller-effect-size transcriptomic signals — genes with modest fold changes are detected in only one or two references depending on exactly where the healthy reference centroid falls — consistent with the higher reference-sensitive gene count for AD in Table 2.

### Non-Lesional Skin in Both Diseases Is Partially Displaced Toward the Lesional Transcriptomic State

To position non-lesional skin along the healthy-to-lesional trajectory, we computed spectrum scores — the scalar projection of each sample’s displacement from the healthy centroid onto the disease axis, expressed as a percentage of the full healthy-to-lesional axis length. Group-level analysis revealed that AD_NL had traversed 22.6% of the healthy-to-AD_LS axis (bootstrap 95% CI 15.2–29.6%), compared with 12.9% for PSO_NL (95% CI 8.4–18.2%; Wilcoxon rank-sum test on individual spectrum scores: W = 2863, p = 0.012; Figure 3B). Cosine similarity confirmed that displacement vectors in both diseases are oriented toward the lesional state (AD cosine=0.772, angle=39.5°; PSO cosine=0.786, angle=38.1°), with near-identical angles (difference 1.4°) indicating shared relative geometric direction but differing magnitude (Figure 3A).

**Figure 3A:**
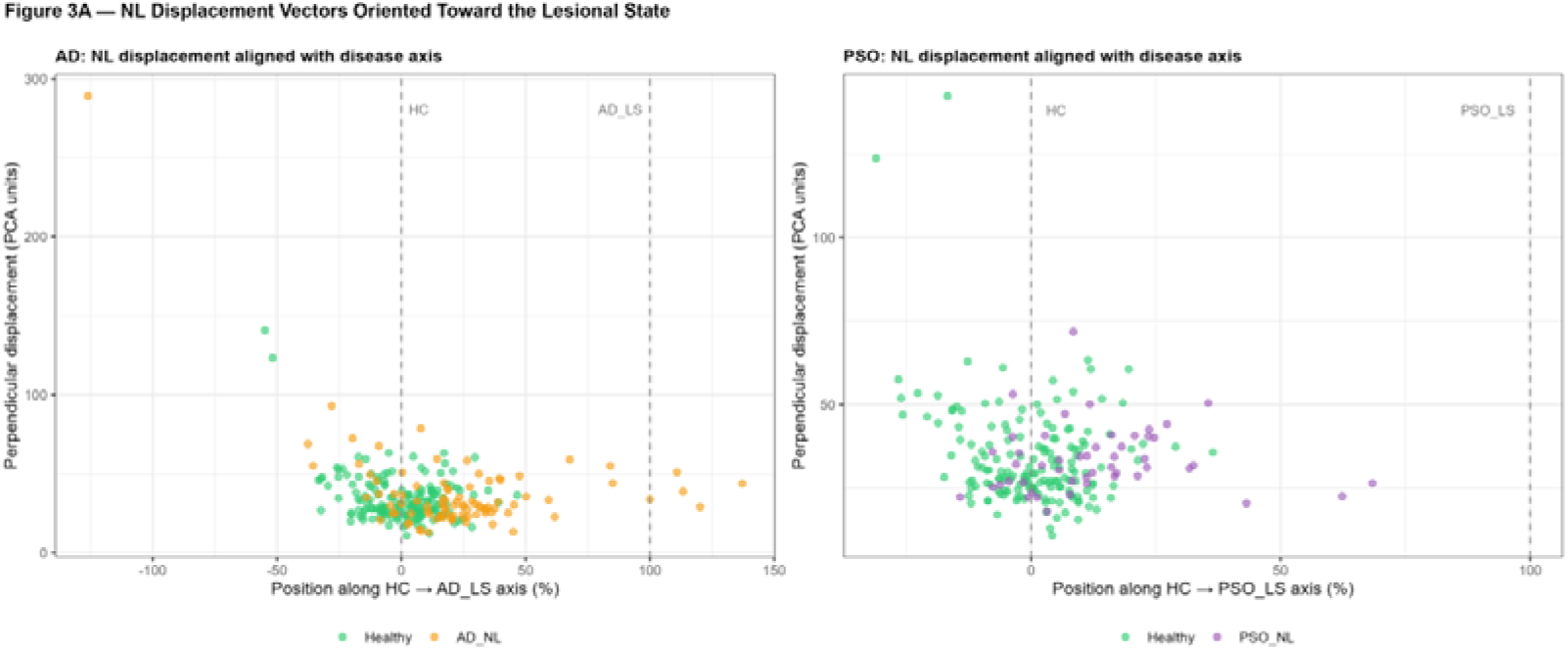
Scatter plots of individual sample positions along (x-axis, spectrum score, % of HC→LS distance) and perpendicular to (y-axis, PCA units) the disease axis, for AD (left, orange = AD_NL, green = Healthy) and PSO (right, purple = PSO_NL, green = Healthy). Dashed vertical lines mark the healthy centroid (0%) and lesional centroid (100%). In the AD panel, AD_NL samples are broadly dispersed, with several exceeding 50% of the disease axis and two samples showing extreme perpendicular displacement (>120 PCA units), indicating movement in a direction orthogonal to the canonical disease trajectory rather than toward or away from the lesional state. In the PSO panel, PSO_NL samples overlap almost entirely with the healthy cluster near 0%, confirming near-healthy transcriptomic positioning. Cosine similarity between non-lesional group displacement and the disease axis: AD = 0.772 (angle 39.5°); PSO = 0.786 (angle 38.1°); Δangle = 1.4°.

**Figure 3B:**
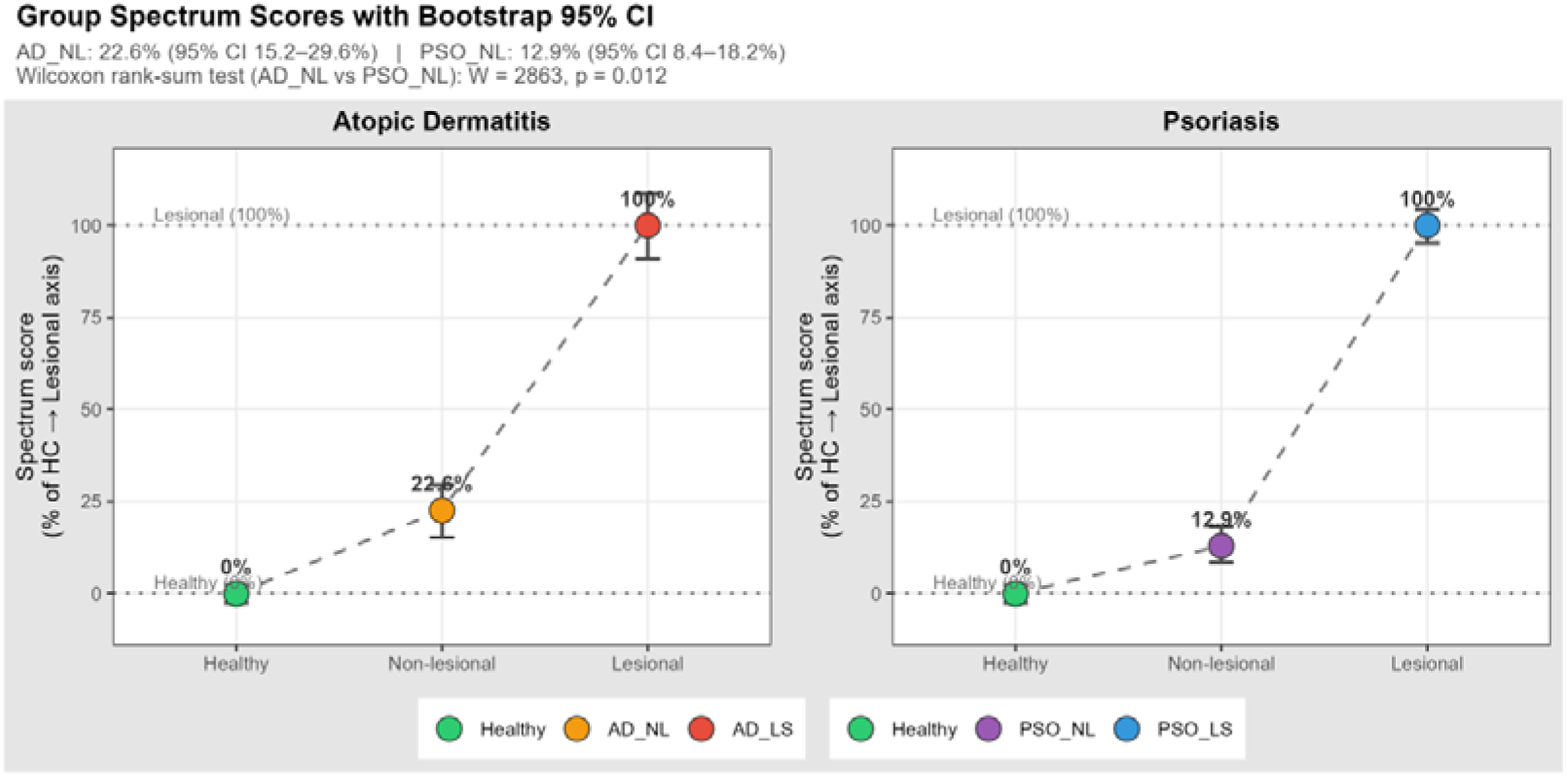
Group-level mean spectrum scores with bootstrap 95% confidence intervals (B = 1,000 resamples, set.seed(42)) along the healthy-to-lesional disease axis for atopic dermatitis (left panel) and psoriasis (right panel). Spectrum score = scalar projection of each sample’s displacement vector from the healthy centroid onto the unit HC→LS disease-axis vector, expressed as a percentage of the full healthy-to-lesional centroid axis length. Each coloured point is a group mean; error bars are bootstrap 95% CIs. Groups shown: Healthy (green), Non-lesional (AD_NL orange / PSO_NL purple), Lesional (AD_LS red / PSO_LS blue). Dashed lines connect the three groups within each disease panel to illustrate the stepwise healthy→non-lesional→lesional progression. Dotted horizontal reference lines mark 0% (healthy centroid) and 100% (lesional centroid) by construction. AD_NL: mean 22.6% (95% CI 15.2–29.6%); PSO_NL: mean 12.9% (95% CI 8.4–18.2%). Wilcoxon rank-sum test on individual spectrum scores confirms a significant group difference (W = 2863, p = 0.012): non-lesional AD skin has traversed a significantly greater fraction of the healthy-to-lesional transcriptomic trajectory than non-lesional psoriasis skin.. All analyses performed in the 20-PC disease-informative coordinate system.

AD_NL scores ranged from −126.3% to +138.2% (values below 0% indicate that a sample’s displacement from the healthy centroid projects opposite to the disease axis; values above 100% indicate projection beyond the lesional centroid along the axis), reflecting the substantial inter-individual variability in the direction and magnitude of transcriptomic departure from homeostasis among AD non-lesional samples. (Figure 3C)(Supplementary Figure S2).

**Figure 3C:**
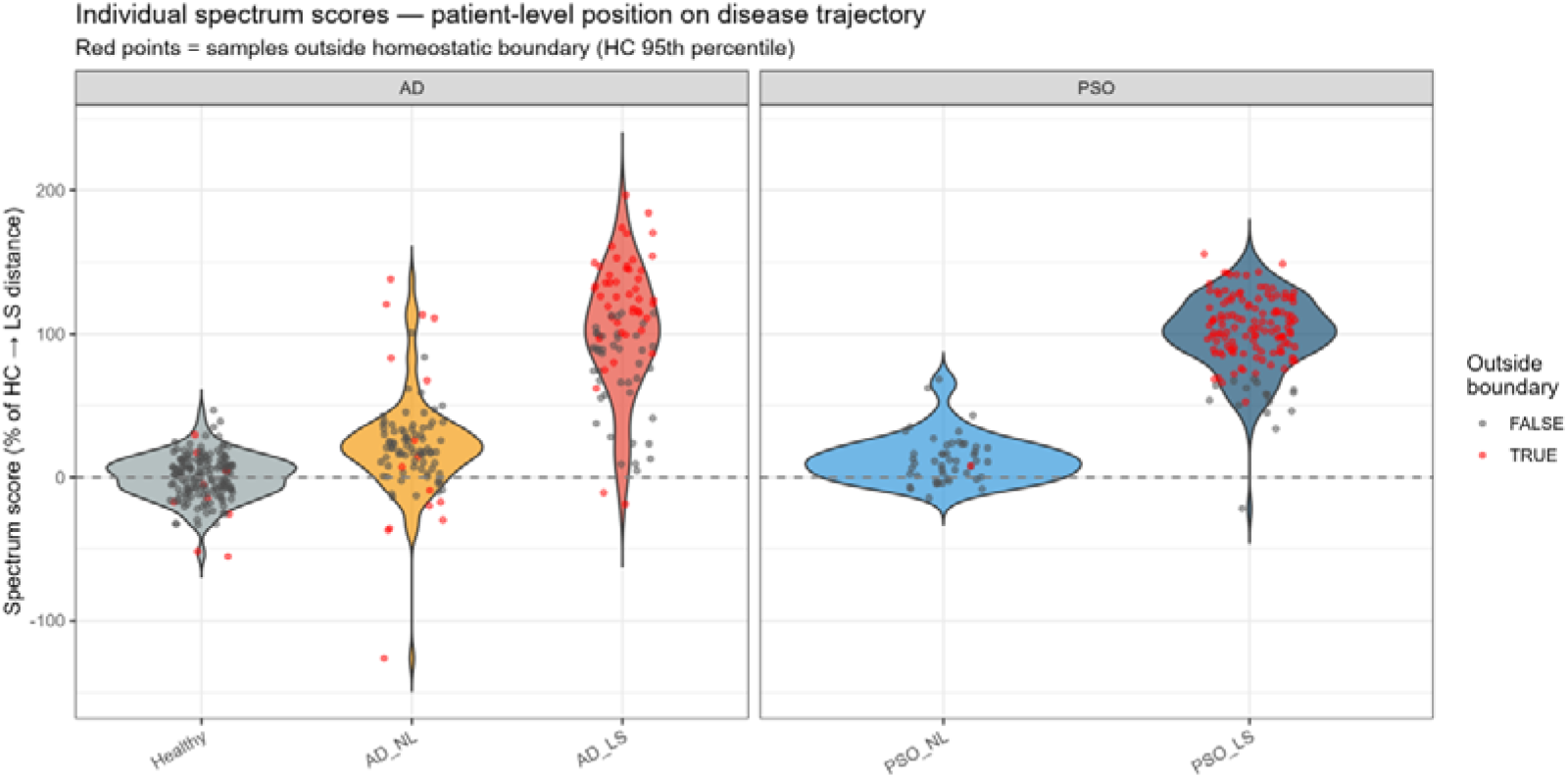
Violin plots with overlaid jittered individual data points showing the full distribution of spectrum scores for all five groups (Healthy, AD_NL, AD_LS, PSO_NL, PSO_LS), split by disease (AD left, PSO right). Red points indicate samples with normalised Euclidean distance from the healthy centroid exceeding the HC 95th-percentile boundary (radius = 53.74 PCA units); grey points are within-boundary. The ‘outside boundary’ designation is determined by Euclidean distance from the healthy centroid, not by spectrum score threshold. AD_NL (orange violin) shows substantially greater distributional spread than PSO_NL (blue violin), with spectrum scores ranging from −126.3% to +138.2%; values below 0% indicate displacement opposite to the disease axis and values above 100% indicate projection beyond the lesional centroid. PSO_NL is tightly confined near 0%. The 2.1-fold greater standard deviation of AD_NL versus PSO_NL (SD 35.0% vs 16.6%) is visible from the violin width. PSO_LS shows the most extreme upward extension, consistent with its high mean normalised distance (1.344).

### A Normalised Distance Gradient Confirms Biological Ordering Independent of Disease Axis

To provide a coordinate-system-independent assessment of group-level displacement, we computed normalised Euclidean distances from the healthy centroid (scaled to the 95th-percentile healthy radius). This analysis depends solely on distances, not axis direction: Healthy 0.616 ≈ PSO_NL 0.634 < AD_NL 0.717 < AD_LS 1.037 << PSO_LS 1.344. PSO non-lesional skin sits virtually atop the healthy cloud (normalised distance 0.634, nearly identical to Healthy at 0.616), whereas AD non-lesional skin is meaningfully further displaced (0.717). This perfect biological gradient — Healthy ≈ PSO_NL < AD_NL < AD_LS << PSO_LS — constitutes robust, axis-independent evidence of disease-specific ordering, aligning with recent multi-cohort observations of AD’s greater non-lesional displacement [10,11].

### AD but Not PSO Non-Lesional Skin Individually Exits the Healthy Homeostatic Transcriptomic Space

The homeostatic boundary was defined as the 95th-percentile Euclidean distance of healthy controls from their centroid (radius=53.74 PCA units). By construction, 5.4% of healthy controls (9/166) exceed this radius. Three independent Fisher exact tests evaluated boundary-crossing significance (Table 3; Figure 4).

**Figure 4:**
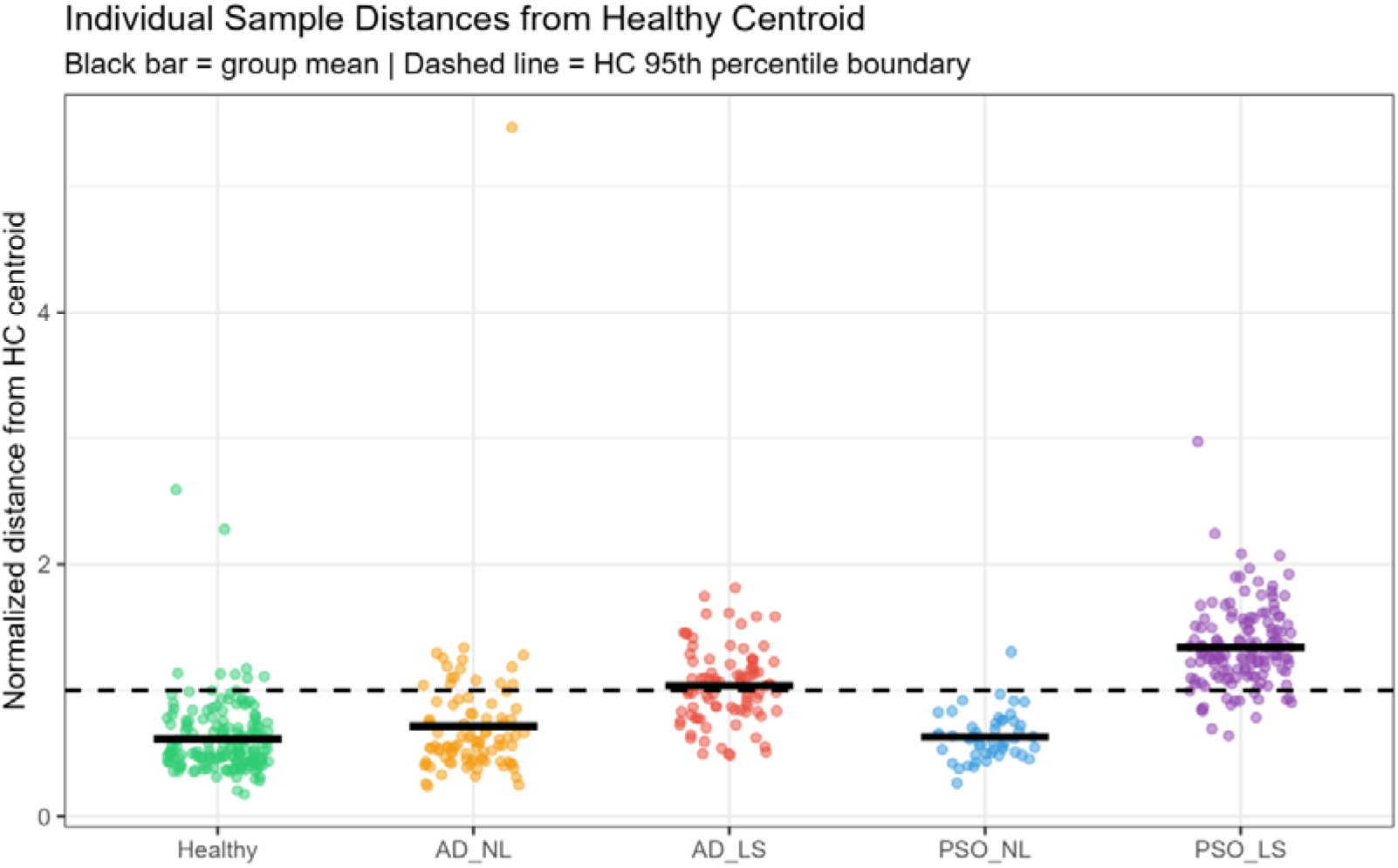
Strip plot of individual normalised Euclidean distances from the healthy centroid for all 537 samples, organised by biological group. Normalised distance = raw Euclidean distance / HC 95th-percentile radius (53.74 PCA units); a value of 1.0 (dashed line) defines the homeostatic boundary. Black horizontal bars indicate group means. The biological gradient is clearly visible: Healthy (green, mean 0.616) and PSO_NL (blue, mean 0.634) cluster predominantly below the boundary; AD_NL (orange, mean 0.717) shows modest upward displacement with 16 of 93 samples (17.2%) crossing the dashed line; AD_LS (red, mean 1.037) sits at or above the boundary; PSO_LS (purple, mean 1.344) lies predominantly above it. The extreme AD_NL outlier (normalised distance ≈5.3) corresponds to GSM5788875 (spectrum score −126.3%), which was confirmed in sensitivity analysis not to drive the boundary-crossing result.

**Table 3:**
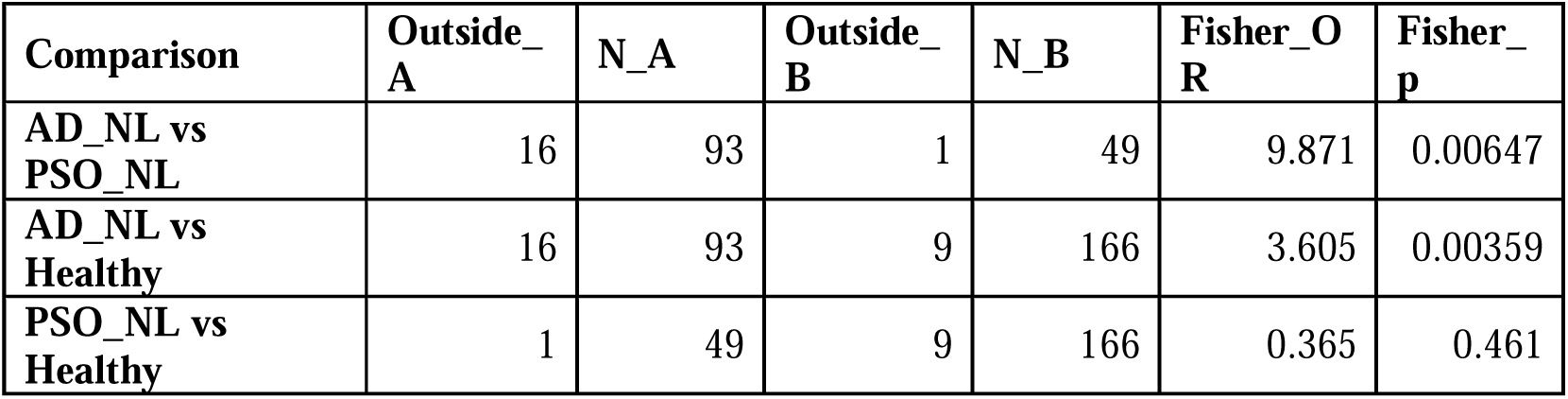
Fisher exact test (mid-P estimate) results for homeostatic boundary-crossing comparisons across three group pairs. AD_NL vs PSO_NL: 16/93 outside vs 1/49 outside, OR = 9.871, p = 0.00647. AD_NL vs Healthy: 16/93 vs 9/166, OR = 3.605, p = 0.00359. PSO_NL vs Healthy: 1/49 vs 9/166, OR = 0.365, p = 0.461. The homeostatic boundary is defined as the 95th-percentile Euclidean distance from the healthy centroid in the 20-PC disease-informative coordinate system (radius = 53.74 PCA units, normalised to 1.0 in Figure 4). All p-values use mid-P Fisher exact estimates as described in the Statistical Analysis section.

PSO_NL showed 1/49 (2.0%) samples outside the boundary (OR=0.36, p=0.461), suggesting that PSO non-lesional skin is transcriptomically indistinguishable from healthy. AD_NL showed 16/93 (17.2%) outside the boundary (OR=3.61, p=0.004), confirming significant displacement. The direct comparison of AD_NL versus PSO_NL yielded OR=9.87 (p=0.007), replicated by permutation testing (n=10,000; p=0.005, observed OR in top 0.5% of null distribution)(Supplementary Figure S3).

This finding replicated consistently across the three cohorts contributing AD_NL samples: GSE121212 (USA) 22.2% (6/27), GSE193309 (Denmark) 11.8% (6/51),

GSE306437 (Ireland) 26.7% (4/15). Across two PSO_NL cohorts, rates were 0/27 (GSE121212) and 1/22 (4.5%; GSE306437).

Robustness analyses confirmed stability: PC sensitivity yielded OR=7.04 (p=0.035) at 10 PCs and OR=9.87 (p=0.007) at 15/20 PCs(supplementary Table S2).

Mahalanobis distance boundary, which accounts for the covariance structure of the healthy reference cloud, confirmed significantly elevated boundary escape in AD_NL versus PSO_NL (AD_NL 52.7% vs PSO_NL 32.7%; OR=2.28, p=0.033)(Supplementary Table S3),

Mahalanobis classification agreed with Euclidean classification in 65% of samples; the smaller OR under Mahalanobis distance reflects boundary-crossing alongdirections of high inter-individual healthy variance — dimensions in which Mahalanobis applies a heavier penalty — a biologically informative rather than artifactual observation, as these are precisely the dimensions driven by disease-enriched genes (CXCL8, FLG2, CCL20, RORC) that are silent in health and activated in AD. the coordinate-system-independent finding is the consistent, robust elevation of AD_NL boundary escape relative to PSO_NL across all distance metrics and cohorts.

### Boundary-Crossing AD Non-Lesional Samples Show Two Directional Patterns with Distinct Pathway Biology

The 16 boundary-crossing AD_NL samples were not uniform. Spectrum scores ranged from −126.3% to +138.2% (mean=25.4%, SD=73.5%), with a natural gap of 41.7 spectrum-score points separating ten samples (≤25.9%) from six samples (≥67.5%; Figure 5A).

**Figure 5A:**
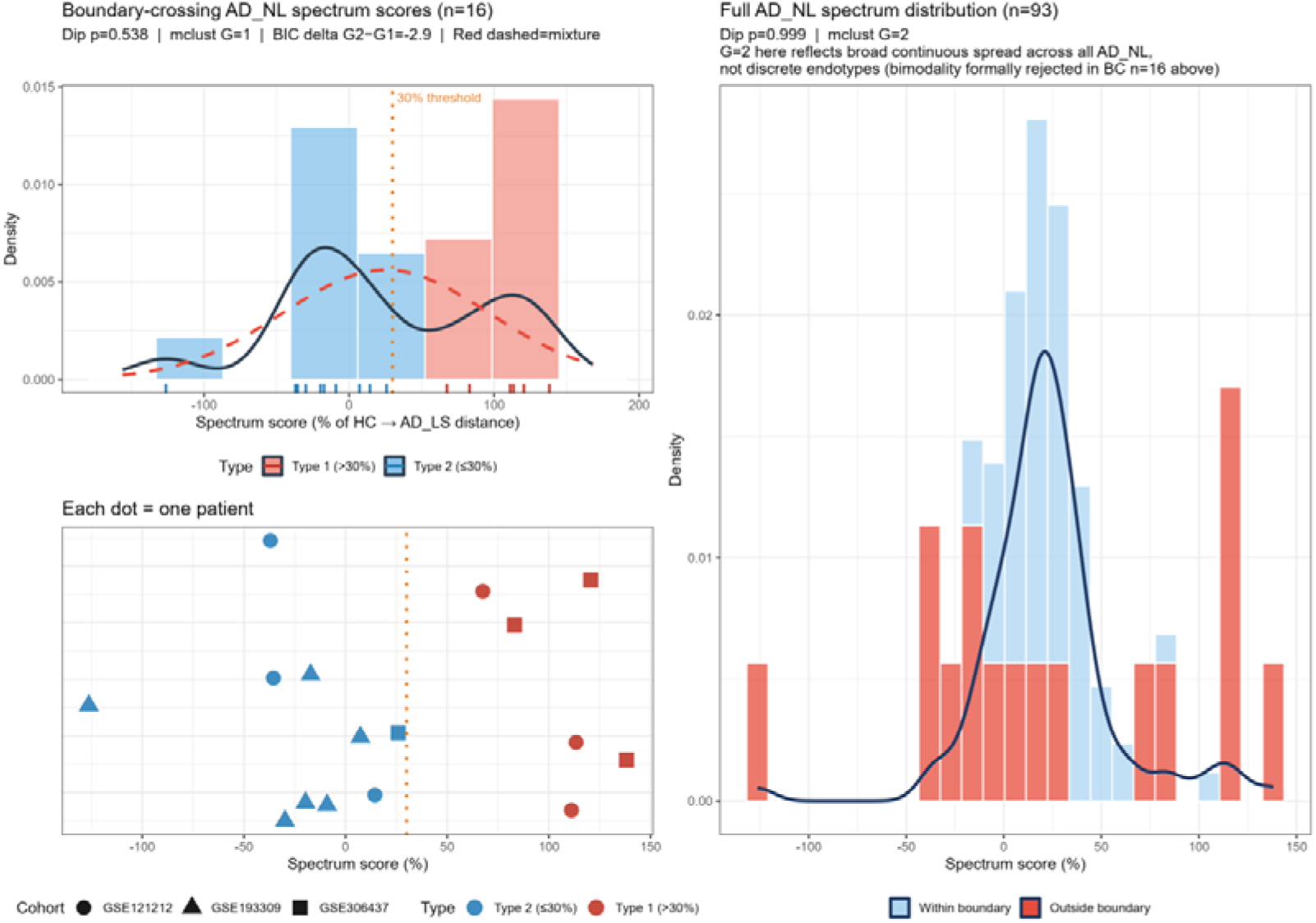
Characterisation of spectrum score distributions among boundary-crossing AD_NL samples (n = 16; left panels) and the full AD_NL cohort (n = 93; right panel). Top-left: density histogram of spectrum scores for n = 16 boundary-crossing samples, coloured by directional pattern: high-positive-displacement Type 1 (red, spectrum > 30%, n = 6) and low/negative-displacement Type 2 (blue, spectrum ≤30%, n = 10). Solid black curve = kernel density estimate; red dashed curve = best-fit single-component Gaussian (mclust G = 1, the selected model). Orange dotted vertical line marks the 30% classification threshold. Formal bimodality tests in this subset were non-significant: Hartigan dip test p = 0.538; mclust BIC selected G = 1 (BIC delta G2−G1 = −2.9, favouring unimodal). The two directional patterns are therefore reported as descriptive observations reflecting continuous heterogeneity, not as validated discrete endotypes. Bottom-left: per-patient strip chart showing individual sample positions, with symbol shape indicating cohort of origin (circles = GSE121212; triangles = GSE193309; squares = GSE306437) and colour indicating directional pattern. Both patterns distribute across at least two independent international cohorts, excluding a study-specific technical artefact. Right panel: spectrum score distribution for the full AD_NL cohort (n = 93), with bars coloured by homeostatic boundary status (light blue = within-boundary n = 77; red = outside boundary n = 16). Dip p = 0.999; mclust selects G = 2 on this full distribution. The G=2 result on the full n=93 distribution is expected by construction: the boundary separates a large within-boundary cluster (n=77, concentrated near 0–30%) from a dispersed boundary-crossing tail (n=16), and mclust detects this imposed separation rather than latent biological discreteness. The appropriate bimodality test — conducted in the boundary-crossing subset where directional heterogeneity is the question — was non-significant by both methods (dip p=0.538, mclust G=1), confirming that the two directional patterns represent continuous heterogeneity within a spectrum rather than validated discrete endotypes.; the formal bimodality assessment was conducted and rejected in the boundary-crossing subset (n = 16, left panels), which constitutes the appropriate test population for directional-pattern heterogeneity.

Formal bimodality testing was underpowered at n=16 (Hartigan dip test p=0.538; mclust BIC delta G2−G1=−2.9, favouring unimodal) and cannot confirm discrete clustering [20,21]. The two directional patterns are therefore reported as observational characterisations, not validated endotypes. The 30% threshold classification was stable across 30–50% thresholds (consistently n=6 high-positive, n=10 low/negative); only at 20% did a single sample shift groups (GSM9201351, spectrum=25.86%), whose displacement is dominated by perpendicular rather than axis-directed movement, supporting its low/negative classification at the primary threshold. To characterise the geometry of each pattern further, displacement vectors for all 16 samples were projected onto both the AD and PSO disease axes. The high-positive group (n=6) showed strong AD-axis alignment (mean AD-axis score 85.4%, cosine 0.44–0.89) with substantial PSO-axis co-projection (mean 59.6%, SD=21.2%), the latter expected given the 30.7° angle between the two disease axes rather than reflecting PSO biology. The low/negative group (n=10), by contrast, showed near-zero alignment with both axes (mean AD-axis score −14.9%, cosine −0.26 to +0.19; mean PSO-axis score −14.9%, cosine −0.30 to +0.12). Residual axis decomposition confirmed that 96.2% of the low/negative group’s boundary-crossing displacement lies outside both disease axes combined (AD axis 2.8%, PSO axis 1.0%, residual 96.2%)(Supplementary Figure S4, Supplementary Table S4)), distributed across three independent cohorts (GSE193309 Denmark n=6, GSE121212 USA n=3, GSE306437 Ireland n=1), ruling out a batch or cohort artefact.

This displacement is carried primarily by PC3 and represents departure along a biological axis orthogonal to both canonical AD and PSO disease trajectories.

High-positive-displacement samples (spectrum >30%, n=6; GSE121212 n=3, GSE306437 n=3) exhibited GSVA pathway scores indicative of systemic inflammatory pre-activation: IL-6/JAK-STAT3 signalling (mean 0.463), allograft rejection (0.387), interferon-alpha response (0.382), inflammatory response (0.379), and TNFα/NF-κB signalling (0.367). This profile aligns with canonical Th2/JAK pre-activation signatures reported in established AD, described here descriptively given n=6 [1,2].

Low/negative-displacement samples (spectrum ≤30%, n=10; GSE121212 n=3, GSE193309 n=6, GSE306437 n=1) showed a markedly different profile: 12 metabolic and stress-response pathways were suppressed relative to within-boundary AD_NL samples at nominal significance (all Wilcoxon p<0.05; BH-adjusted values 0.052–0.18, FDR-trending; Table 4, Figure 5B).

**Figure 5B:**
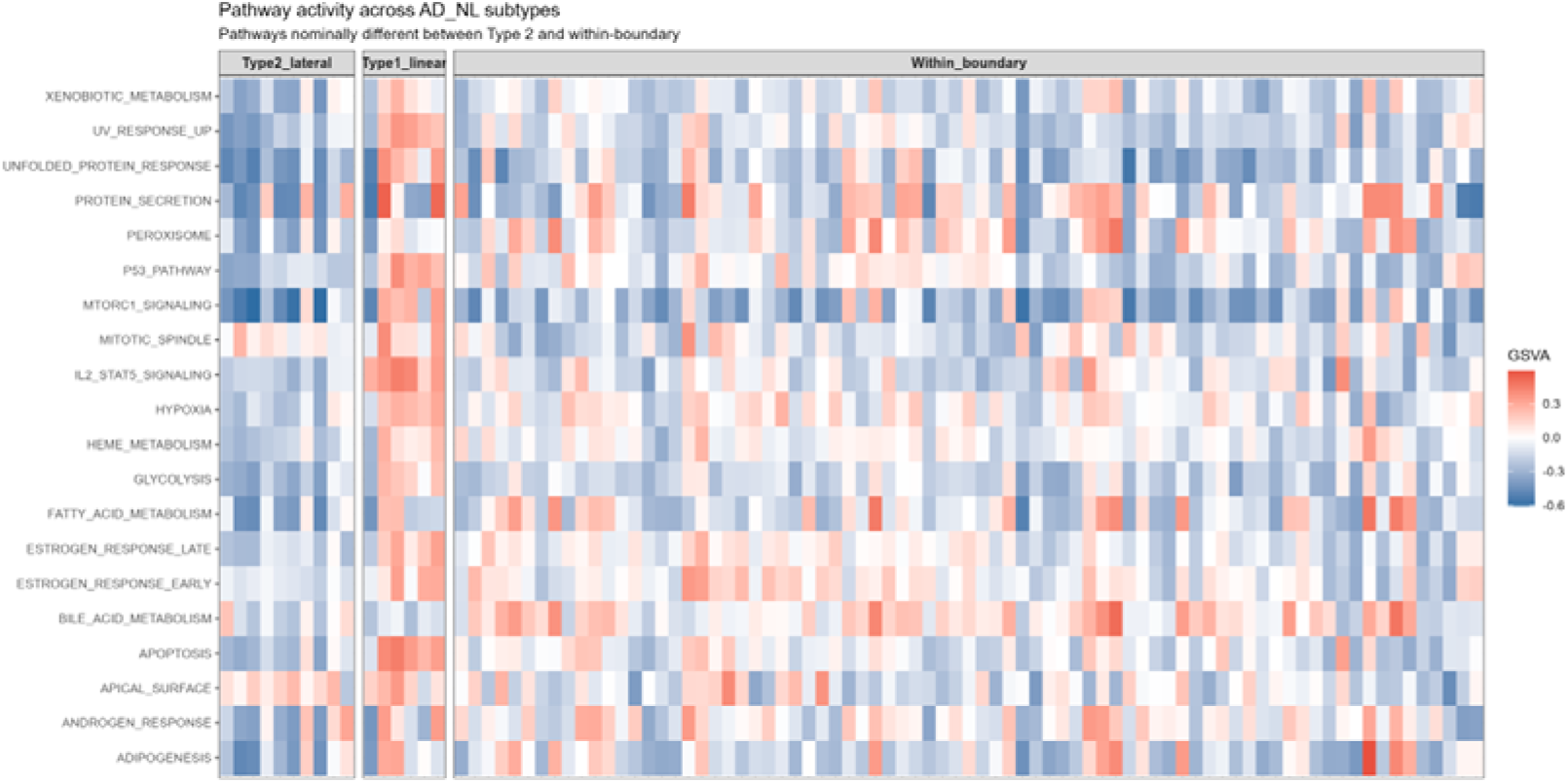
Heatmap of GSVA pathway enrichment scores (MSigDB Hallmark gene sets, n = 50, kcdf = ‘Gaussian’) across AD_NL subgroups: low/negative-displacement (Type 2 lateral, n = 10; left columns), high-positive-displacement (Type 1 linear, n = 6; middle columns), and within-boundary AD_NL (n = 77; right columns). Each column is one sample; the 20 pathways showing nominal Wilcoxon difference between Type 2 and within-boundary samples (p < 0.05 uncorrected) are shown as rows. Colour scale: red = positive GSVA enrichment score, blue = negative (pathway suppressed), white = near-zero. The low/negative-displacement group (Type 2, left) shows consistent deep blue across metabolic and stress-response pathways (MTORC1_SIGNALING, HEME_METABOLISM, HYPOXIA, UNFOLDED_PROTEIN_RESPONSE, GLYCOLYSIS, FATTY_ACID_METABOLISM, P53_PATHWAY), with APICAL_SURFACE paradoxically elevated (red). The high-positive-displacement group (Type 1, middle) shows mixed warmer tones, consistent with inflammatory pre-activation. The two groups are visually discriminable at the pathway level despite formal bimodality tests not reaching significance, suggesting functionally distinct transcriptional programs within the boundary-crossing population.

**Table 4:** Wilcoxon rank-sum test results for GSVA Hallmark pathway activity differences between the low/negative-displacement boundary-crossing subgroup (n = 10) and within-boundary AD_NL samples (n = 77). Twelve pathways showing nominal significance (Wilcoxon p < 0.05) are listed, with Benjamini–Hochberg adjusted p-values. No pathway reaches the FDR threshold of 0.05 after correction (range: BH-adjusted p = 0.052–0.177). Results should be interpreted as hypothesis-generating directional signals. Most suppressed pathways in the low/negative group: Heme Metabolism (p = 0.0014, BH = 0.052), P53 Pathway (p = 0.0043, BH = 0.052), Estrogen Response Late (p = 0.0047, BH = 0.052), Hypoxia (p = 0.0062, BH = 0.052), Unfolded Protein Response (p = 0.0065, BH = 0.052).

| Pathway | Mean GSVA<br>(Low/negative<br>n=10) | Mean<br>GSVA<br>(Within-<br>boundar<br>y n=77) | Differenc<br>e<br>(Low/neg<br>minus<br>Within) | P-value<br>(Wilcoxon<br>) | P-adj<br>(BH) |
| --- | --- | --- | --- | --- | --- |
| Heme Metabolism | -0.17 | -0.032 | -0.138 | 0.00137 | 0.0524 |
| P53 Pathway | -0.208 | -0.061 | -0.147 | 0.00431 | 0.0524 |
| Estrogen Response Late | -0.16 | -0.031 | -0.129 | 0.00469 | 0.0524 |
| Hypoxia | -0.15 | -0.006 | -0.143 | 0.00624 | 0.0524 |
| Unfolded Protein | -0.348 | -0.174 | -0.173 | 0.0065 | 0.0524 |

| Response |  |  |  |  |  |
| --- | --- | --- | --- | --- | --- |
| Peroxisome | -0.206 | -0.007 | -0.199 | 0.0112 | 0.0649 |
| Apoptosis | -0.189 | -0.055 | -0.134 | 0.0117 | 0.0649 |
| Bile Acid Metabolism | -0.065 | 0.081 | -0.147 | 0.0209 | 0.105 |
| Glycolysis | -0.248 | -0.137 | -0.111 | 0.0233 | 0.106 |
| Estrogen Response Early | -0.09 | 0.001 | -0.09 | 0.0276 | 0.114 |
| Uv Response Up | -0.236 | -0.112 | -0.124 | 0.0296 | 0.114 |
| Fatty Acid Metabolism | -0.228 | -0.049 | -0.179 | 0.0497 | 0.177 |

12 Suppressed pathways included heme metabolism (p=0.0014), p53 pathway (p=0.0031), hypoxia (p=0.0044), unfolded protein response (p=0.0061), mTORC1 signalling (p=0.0088), peroxisome biogenesis (p=0.0092), fatty acid metabolism (p=0.0104), and glycolysis (p=0.0129). Paradoxically, the apical surface pathway (tight junction and epithelial barrier markers) was elevated (p=0.0038). Critically, IFN-alpha response (mean GSVA −0.260), IFN-gamma response (−0.240), and inflammatory response (−0.142) were all suppressed in the low/negative group relative to the high-positive group (+0.382, +0.358, +0.379 respectively), confirming that this pattern does not represent attenuated inflammatory activation but a qualitatively distinct biology. All 12 suppressions remained significant after removing the most extreme sample (GSM5788875, spectrum=−126.3%), with three additional pathways gaining significance. Both directional patterns distributed across at least two independent cohorts, excluding technical artefact.

Cell type deconvolution using EPIC addressed two questions. First, comparing all boundary-crossing AD_NL samples (n=16) versus within-boundary AD_NL (n=77), CD8+ T cell proportions were modestly but significantly elevated in boundary-crossing samples (mean 0.1080 vs 0.1037; Wilcoxon p=0.003), suggesting a small increase in CD8+ infiltrate associated with boundary escape overall [24]. The absolute difference is small (0.004 proportion units) and these estimates carry inherent deconvolution uncertainty, but the signal is consistent with tissue-resident memory T cell roles in maintaining subclinical inflammatory setpoints [7,24]. Second, a three-way comparison of low/negative-displacement (n=10), high-positive-displacement (n=6), and within-boundary AD_NL (n=77) showed no significant differences in any cell type proportion across groups (all Wilcoxon p>0.10; Table 5) — an expected finding given that splitting n=16 into n=6 and n=10 removes statistical power to detect the modest CD8+ signal observed in the combined comparison; both directional subgroups carry this elevation approximately equally rather than one group driving it:

**Table 5:** EPIC immune cell deconvolution results comparing within-boundary AD_NL (n = 77) versus all boundary-crossing AD_NL (n = 16). Wilcoxon rank-sum test, no correction for multiple comparisons across seven cell types. CD8+ T cells: 0.1037 (inside) vs 0.1080 (outside), fold change 1.04, p = 0.0029 (significant at nominal threshold). Uncharacterised cells: 0.6077 vs 0.5977, fold change 0.98, p = 0.014 (significant at nominal threshold). CD4+ T cells (p = 0.076), neutrophils (p = 0.107), monocytes (p = 0.131), NK cells (p = 0.379), B cells (p = 0.923): all non-significant. If Bonferroni correction were applied across seven cell types (threshold p < 0.007), only CD8+ T cells would remain significant.

| Cell_type | Mean_inside | Mean_outside | Fold_change | p_value | Significant |
| --- | --- | --- | --- | --- | --- |
| T cell CD8+ | 0.1037 | 0.108 | 1.04 | 0.0029 | TRUE |
| uncharacterized cell | 0.6077 | 0.5977 | 0.98 | 0.014 | TRUE |
| T cell CD4+ | 0.1083 | 0.1105 | 1.02 | 0.0757 | FALSE |
| Neutrophil | 0.1013 | 0.1044 | 1.03 | 0.1067 | FALSE |
| Monocyte | 0.0222 | 0.0227 | 1.02 | 0.1306 | FALSE |
| NK cell | 0 | 0 | 0 | 0.3786 | FALSE |
| B cell | 0.0567 | 0.0567 | 1 | 0.923 | FALSE |

CD4+ T cells: low/neg 0.108, high-pos 0.114, within-boundary 0.108; CD8+ T cells: low/neg 0.106, high-pos 0.112, within-boundary 0.104; B cells: 0.056, 0.058, 0.057; monocytes: 0.022, 0.024, 0.022. The two directional subgroups share the modest CD8+ elevation associated with boundary-crossing equally — the global metabolic suppression and PC3 displacement in the low/negative group is therefore not attributable to differential cellular composition between subgroups, and the pathway differences between the two patterns reflect transcriptional biology rather than differences in tissue cellular infiltrate. The biological basis of this PC3-axis suppression pattern — whether representing an integrated stress response, a distinct keratinocyte biology, or a clinically heterogeneous subgroup — cannot be determined from bulk RNA-seq alone and requires cell-type-resolved follow-up investigation.

Also, anatomic region (arm vs foot) did not differ between boundary-crossing and within-boundary AD_NL samples (Fisher p=0.594), excluding sampling site as a confound.

## Discussion

The reference sensitivity finding resolves a decade of conflicting AD transcriptomic reports as a geometric inevitability rather than a failure of individual studies. When a disease’s signature depends on genes whose effect sizes cluster near the detection threshold, small shifts in reference composition — reflecting genuine inter-individual variation in healthy skin immune tone, microbiome exposure, and barrier expression — are sufficient to move marginal genes across significance boundaries [12,13]. This is precisely the architecture of AD: reference-sensitive genes are enriched for GTPase signalling and epithelial-mesenchymal transition pathways that show heterogeneous, low-amplitude activity, while reference-stable genes align with epidermal barrier function where signal-to-noise is high [5,9,10]. PSO signatures, anchored in high-amplitude IL-17/keratinocyte proliferation programs, are structurally less vulnerable to this effect [5]. The implication is methodological and reproducibility-relevant: improving cross-study agreement in AD transcriptomics requires either larger healthy reference cohorts to stabilise centroid estimates, or a addtion to difference-based DEG analysis a position-based geometric frameworks that are reference-composition-independent by design.

The spectrum scoring and distance gradient analyses together establish that non-lesional skin is not a neutral baseline in either disease, but the degree of departure differs fundamentally between AD and PSO. PSO non-lesional skin sits virtually coincident with the healthy cloud by both axis-independent (normalised distance 0.634 versus healthy 0.616) and axis-dependent (12.9% spectrum displacement) measures, consistent with PSO’s proposed switch-like transcriptomic transition in which lesional disease activates abruptly from a near-normal non-lesional substrate [5,10,11]. AD non-lesional skin, by contrast, has already traversed 22.6% of the healthy-to-lesional axis and sits measurably further from the healthy centroid (0.717), supporting a gradual subclinical priming model in which immune and barrier dysregulation accumulates continuously before clinical manifestation [5,10,11]. Within each disease, non-lesional displacement is directed toward rather than orthogonal to its own lesional pole (cosine similarity: AD 0.772, 39.5°; PSO 0.786, 38.1°; within-disease Δangle 1.4°) — confirming that both NL states are geometrically primed in the disease direction; the AD and PSO disease axes themselves are 30.7° apart and not coincident, so the near-identical cosines reflect a shared within-disease directional relationship, not a shared biological trajectory. The distinction between AD and PSO non-lesional skin is therefore one of displacement magnitude along divergent disease axes (Wilcoxon W=2863, p=0.012), a separation that is invisible to any analysis that collapses non-lesional skin into a single reference category.

The homeostatic boundary analysis extends the group-level finding to the individual level, revealing that the AD non-lesional group mean conceals a bimodal clinical reality: the majority of AD non-lesional samples (82.8%) remain within the homeostatic space, while a reproducible minority (17.2%, replicated across three independent international cohorts) have individually exited it. The gene programs driving this escape — CXCL8, FLG2, CCL20, RORC — are established mediators of chemokine signalling, barrier compromise, and Th17/Th22 activation that are biologically silent in healthy skin and activated in lesional disease [5,10], suggesting that boundary-crossing samples have initiated a disease-associated transcriptional programme prior to clinical lesion formation. The coordinate-system independence of this finding — confirmed by Mahalanobis distance sensitivity analysis (OR=2.28, p=0.033) and stable across 10–20 PC choices — excludes a geometric artefact explanation. That no comparable signal is seen in PSO non-lesional skin (2.0% boundary-crossing; OR=0.36 vs healthy, p=0.461) aligns with the clinical observation that PSO non-lesional skin is macroscopically and histologically near-normal, while AD non-lesional skin shows subclinical barrier and immune abnormalities even in unaffected sites [3,4,7].

Among boundary-crossing AD non-lesional samples, the two directional patterns are the most novel and open finding of this analysis. The high-positive displacement pattern (n=6) is mechanistically interpretable: IL-6/JAK-STAT3, interferon-alpha, and TNFα/NF-κB pre-activation aligns with canonical Th2/JAK signatures reported in established AD [1,2], and the therapeutic implication — that this subset may be a candidate for early JAK inhibitor or biologic intervention — is hypothesis-generating and consistent with emerging endotype literature, though it requires prospective validation at this sample size [12,13]. The low/negative displacement pattern (n=10) is more intriguing and less tractable. With 96.2% of displacement carried by a biological axis orthogonal to both the AD and PSO disease trajectories, global suppression of 12 metabolic and stress-response pathways (heme metabolism, mTORC1, hypoxia, unfolded protein response, glycolysis, fatty acid metabolism), and paradoxical elevation of apical surface/barrier markers, this pattern does not represent attenuated inflammatory activation — it represents a qualitatively different biology. Cell type deconvolution confirms it is not driven by differential cellular infiltration; both directional subgroups share the modest CD8+ elevation associated with boundary-crossing equally (three-way comparison all p>0.10), and the metabolic suppression profile therefore reflects transcriptional rather than compositional differences. Whether this pattern represents keratinocyte integrated stress response, filaggrin-null intrinsic metabolic failure, a clinically heterogeneous subgroup selected by shared geometric position rather than shared mechanism, or an upstream biology preceding canonical inflammatory activation, cannot be resolved from bulk RNA-seq alone. It constitutes a testable hypothesis for spatial transcriptomics or single-cell investigation of non-lesional AD skin.

The modest but statistically significant CD8+ T cell elevation in boundary-crossing versus within-boundary AD non-lesional samples (mean 0.1080 vs 0.1037; p=0.003) is consistent with a tissue-resident memory T cell role in maintaining subclinical inflammatory setpoints in AD non-lesional skin [7,24]. The absolute difference is small (fold change 1.04) and carries inherent deconvolution uncertainty, but it points toward a cellular correlate of geometric boundary escape that warrants orthogonal validation by spatial transcriptomics or immunohistochemistry. Crucially, this elevation is shared equally across both directional subgroups, reinforcing that the pathway biology distinguishing the two patterns is transcriptional in origin.

## Conclusion

This study establishes three findings in sequence. First, AD transcriptomic inconsistency across studies is a structural consequence of AD’s reliance on smaller-effect-size genes, not a failure of methodology — a mechanistic explanation that points toward reference-independent geometric positioning as a more reproducible analytical framework. Second, non-lesional skin is not a uniform baseline in either disease: AD non-lesional skin occupies a measurably disease-proximal position along the healthy-to-lesional transcriptomic axis, while PSO non-lesional skin remains near-healthy — a group-level difference that is invisible to analyses that treat non-lesional skin as a single reference category. Third, at the individual level, a reproducible subset of AD non-lesional patients has already exited the homeostatic transcriptomic space entirely, in two biologically distinct directions: one consistent with canonical inflammatory pre-activation, the other pointing toward a third biological axis orthogonal to both known disease trajectories and not attributable to tissue cellular composition. The immediate clinical implication is a reframing of non-lesional skin from a uniform baseline to a heterogeneous landscape in which individual patients occupy very different positions relative to homeostasis before any clinical lesion appears. Whether this geometric positioning predicts lesional onset, treatment response, or disease course requires prospective longitudinal validation — but the framework for asking that question at the individual level, scalable to multi-omics data and applicable to other heterogeneous inflammatory diseases, is now demonstrated.

## Limitations

Several limitations must be acknowledged. First, the analysis is entirely computational and lacks direct functional or histological validation of the boundary-crossing phenotype in patient tissue. Second, immune cell proportion estimates derive from bulk RNA-seq deconvolution using EPIC, which carries inherent uncertainty due to reliance on reference profiles and cannot resolve spatial heterogeneity or distinguish tissue-resident from infiltrating populations [24]. The CD8+ elevation in boundary-crossing samples overall (p=0.003) and the null result across directional subgroups (all p>0.10) are internally consistent but require orthogonal validation via spatial transcriptomics or immunohistochemistry to confirm cell-type-level biology. Third, PSO non-lesional data are available from only two cohorts (n=49 total), providing less statistical power than AD non-lesional samples across three cohorts (n=93), which may influence comparative robustness [10,11]. Fourth, the two directional patterns among boundary-crossing AD non-lesional samples rely on a data-driven spectrum-score threshold without external pre-specification; while stable across 20–50% thresholds and distributed across cohorts, they represent observational characterisations rather than validated discrete endotypes, and prospective studies with larger sample sizes are needed to distinguish true bimodality from a continuous distribution [20,21]. Finally, although ComBat batch correction with a protection matrix is a well-established approach for multi-cohort integration, residual batch effects cannot be completely excluded given the asymmetric distribution of groups across datasets [17].

## Declarations

### Ethics approval and consent to participate

Not applicable. This study was based exclusively on analysis of publicly available transcriptomic datasets from the Gene Expression Omnibus (GEO). No new human or animal samples were collected.

### Availability of data and materials

The datasets analyzed during the current study are openly available in the Gene Expression Omnibus (GEO) repository under the following accession numbers: (GSE121212, GSE193309, GSE306437, GSE54456)

All processed data and analysis scripts are available from the corresponding author upon reasonable request.

---

**Supplementary Figure S1:**
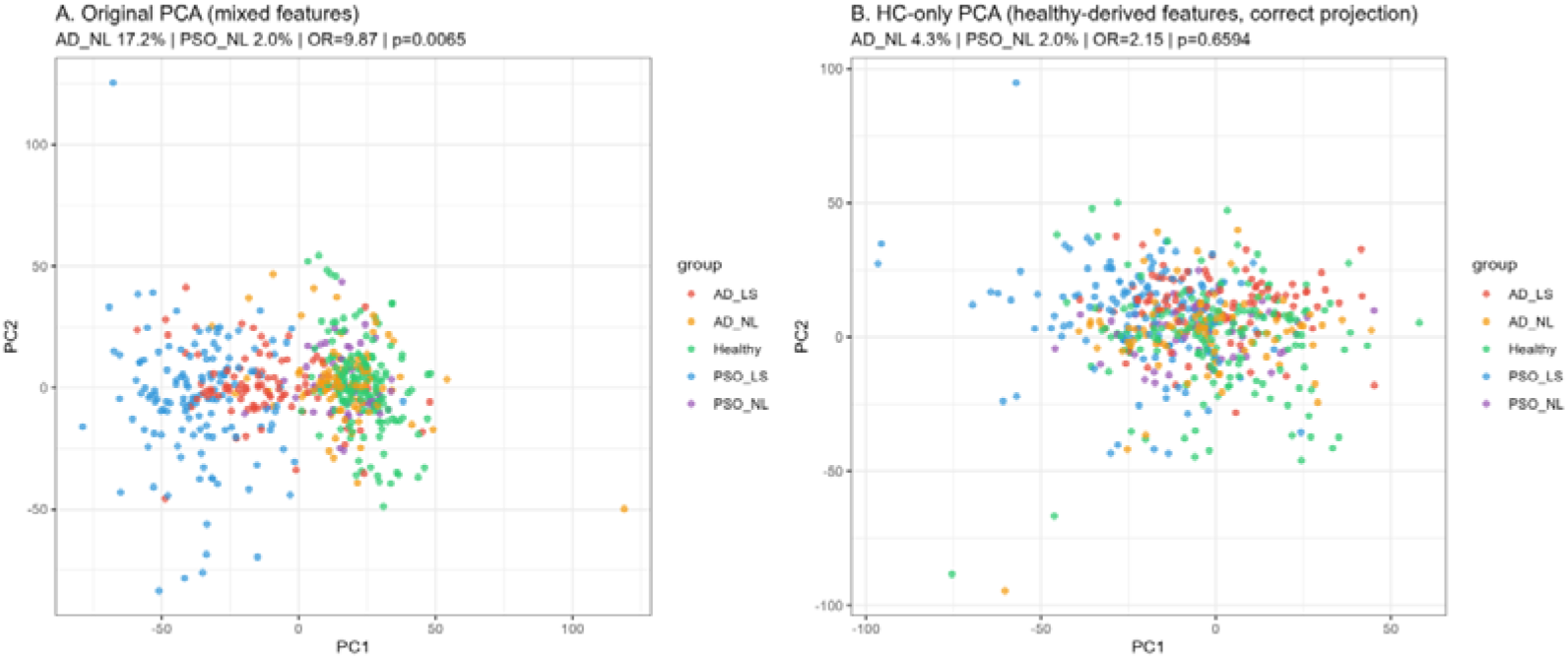
Sensitivity analysis comparing homeostatic boundary results under two coordinate systems. Panel A (left): Primary mixed-feature PCA space, in which the feature set comprises the top 3,000 most variable genes across all 537 samples combined (disease-informative). AD_NL boundary-crossing: 17.2% (16/93); PSO_NL: 2.0% (1/49); OR = 9.87, p = 0.0065. Five biological groups are clearly differentiated. Panel B (right): Healthy-control-only PCA space, in which the feature set is derived from variance in healthy controls alone. Under this coordinate system, all five biological groups (including AD_LS and PSO_LS) collapse into a single overlapping cloud with no group separation, and AD_NL boundary-crossing drops to 4.3% (OR = 2.15, p = 0.659, non-significant). This demonstrates empirically that the disease-informative mixed-feature coordinate system is necessary to resolve the subclinical dysregulation present in non-lesional AD; the healthy-only axes are directed toward healthy variation noise rather than disease signal. The figure provides the empirical motivation for using mixed-feature PCA as the primary coordinate system, as described in the Methods.

**Supplementary Figure S2.**
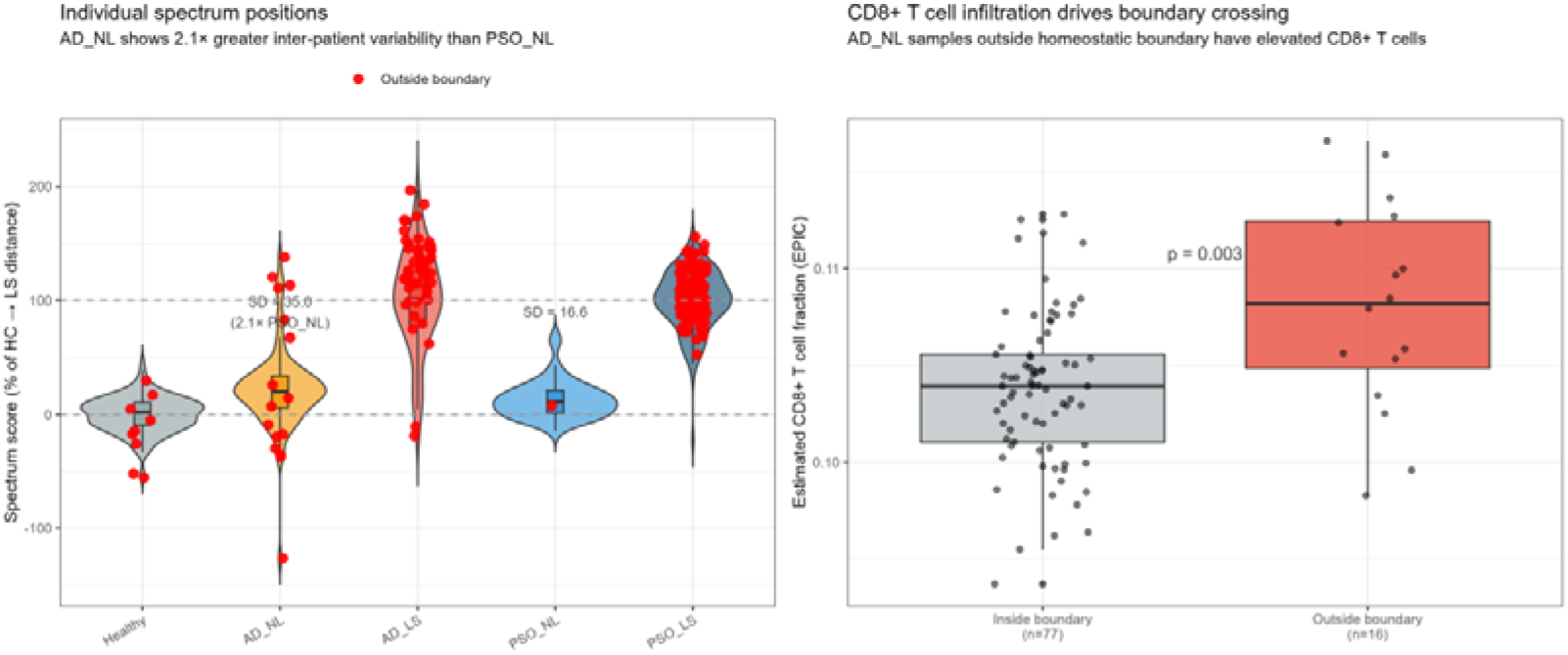
: Two-panel supplementary figure. Left: Violin plots of individual spectrum scores for all five groups (Healthy, AD_NL, AD_LS, PSO_NL, PSO_LS) replicating Figure 3C layout, with SD annotations for the non-lesional groups (AD_NL SD = 35.0%, 2.1× PSO_NL; PSO_NL SD = 16.6%). Red dots indicate boundary-crossing samples; yellow squares indicate group means. Right: Boxplot with individual data points showing estimated CD8+ T cell fraction (EPIC deconvolution algorithm, immunedeconv R package v2.1.0) in within-boundary AD_NL (grey, n = 77) versus boundary-crossing AD_NL (red, n = 16). Boundary-crossing samples show a modest but statistically significant elevation in CD8+ T cell fraction (Wilcoxon rank-sum p = 0.003; mean 0.1080 versus 0.1037 within-boundary). The absolute difference (0.004 proportion units, fold change 1.04) is small, and the distributions overlap substantially; this represents a statistically detectable but biologically modest associative signal.

**Supplementary Figure S3:**
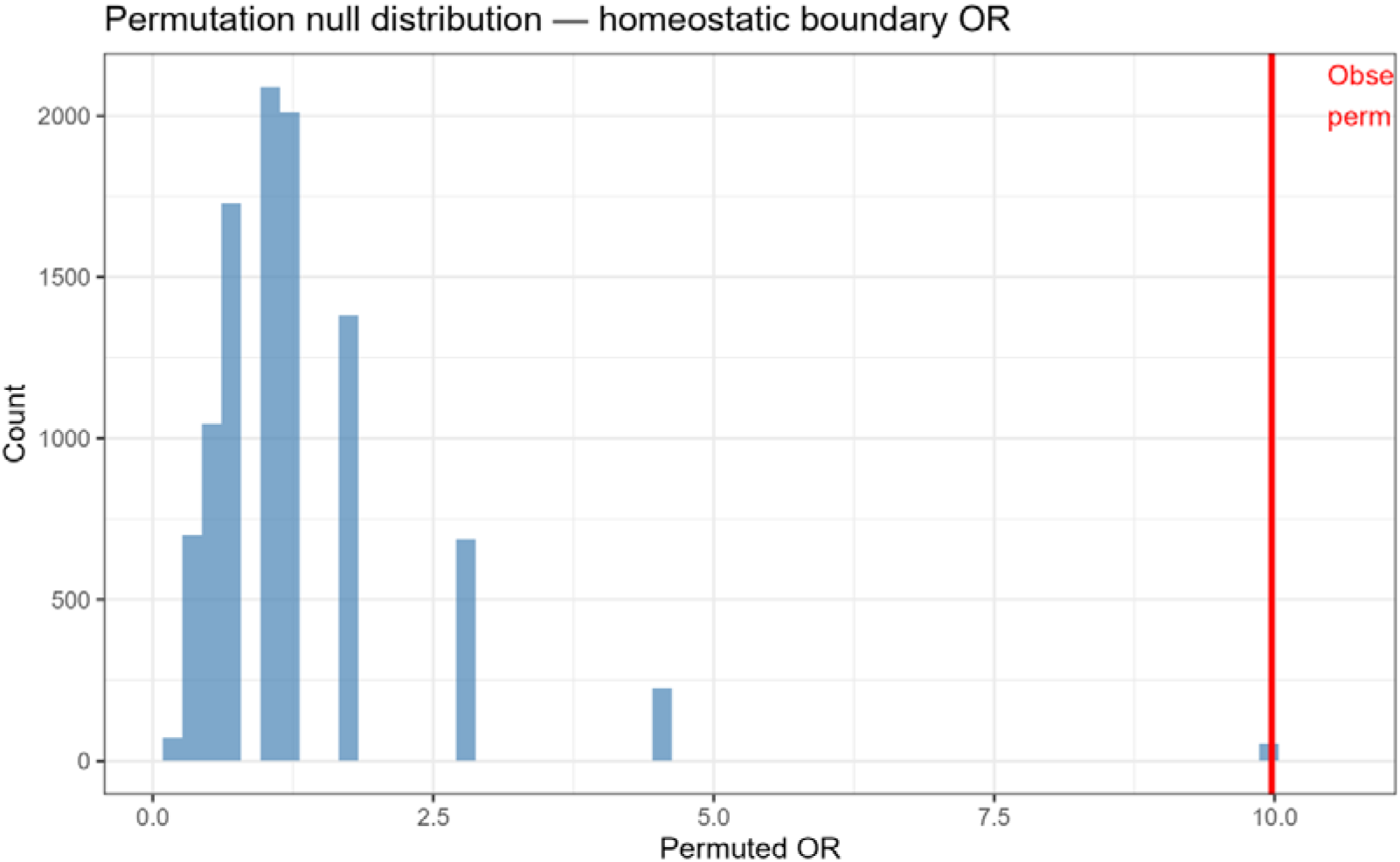
Permutation null distribution for the homeostatic boundary-crossing odds ratio. Histogram shows the distribution of ORs from 10,000 random permutations of AD_NL and PSO_NL group labels (keeping group sizes fixed at n = 93 and n = 49). The null distribution is right-skewed and concentrated between OR = 0 and approximately OR = 2.5. The red vertical line marks the observed OR = 9.87. The empirical permutation p-value is 0.005 (observed OR exceeded by only ≈50 of 10,000 permuted ORs), confirming that the boundary-crossing disparity between AD_NL and PSO_NL cannot be explained by chance label assignment.

**Supplementary Figure S4:**
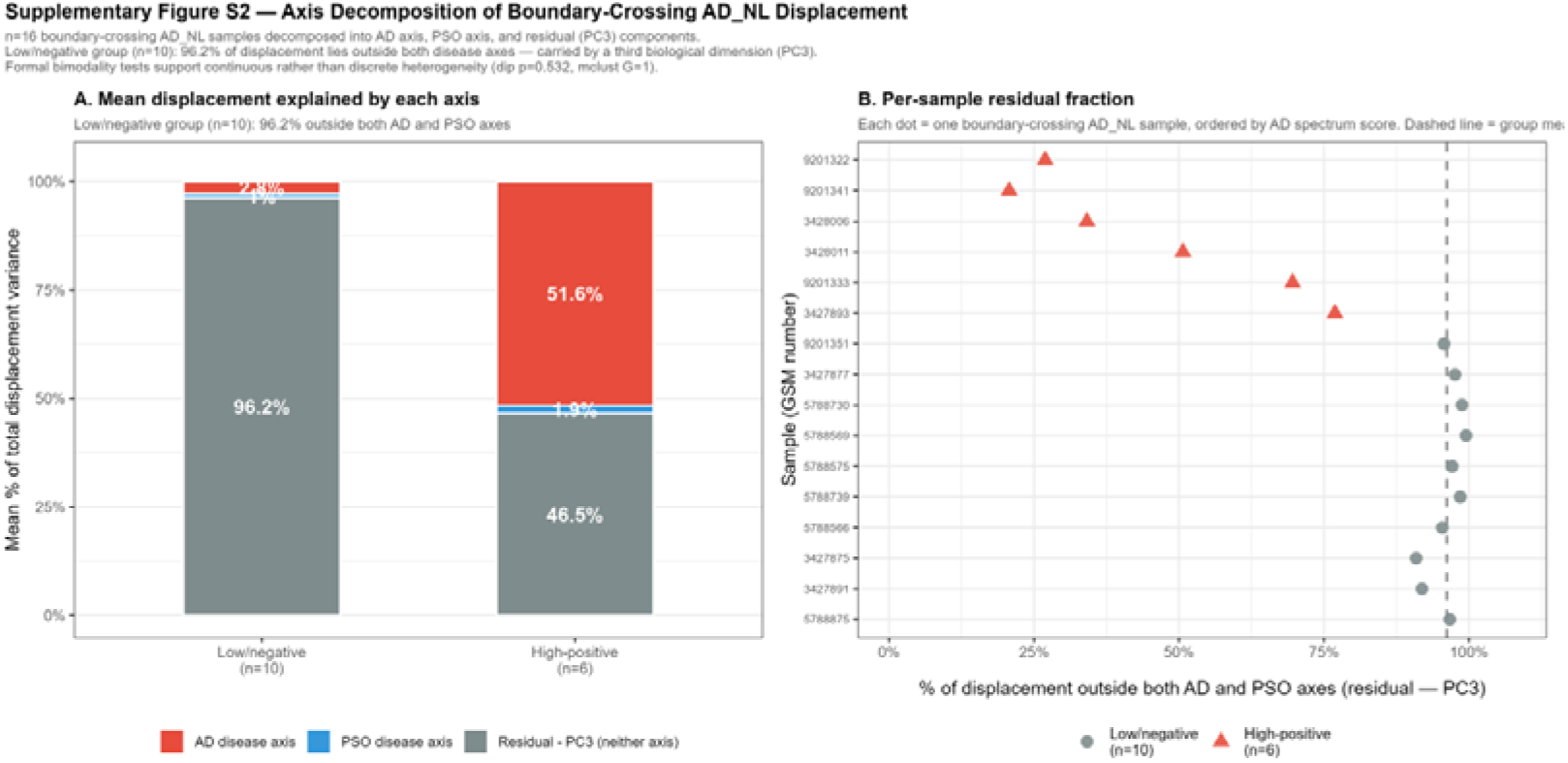
Axis decomposition of displacement vectors for the 16 boundary-crossing AD_NL samples. Panel A (left): Stacked bar chart showing the mean percentage of total displacement variance attributed to the AD disease axis (red), PSO disease axis (blue), and residual third-axis component (grey = PC3 and orthogonal dimensions) for low/negative-displacement (n = 10) and high-positive-displacement (n = 6) groups. Low/negative group: 96.2% residual, 2.3% AD axis, 1.9% PSO axis — boundary-crossing is carried almost entirely by a biological dimension orthogonal to both canonical disease trajectories. High-positive group: 51.6% AD axis, 46.5% residual, 1.9% PSO axis — displacement is primarily aligned with the AD disease direction. Panel B (right): Per-sample dot plot ordered by spectrum score (ascending), showing each sample’s residual fraction (% of displacement outside both AD and PSO axes). Low/negative samples (grey circles) cluster near 91.9–99.5% residual; high-positive samples (red triangles) range from 20.7–76.9% residual. The clear separation of the two groups on the residual axis confirms that their mechanistic basis is structurally distinct.

**Supplementary Table S2:**
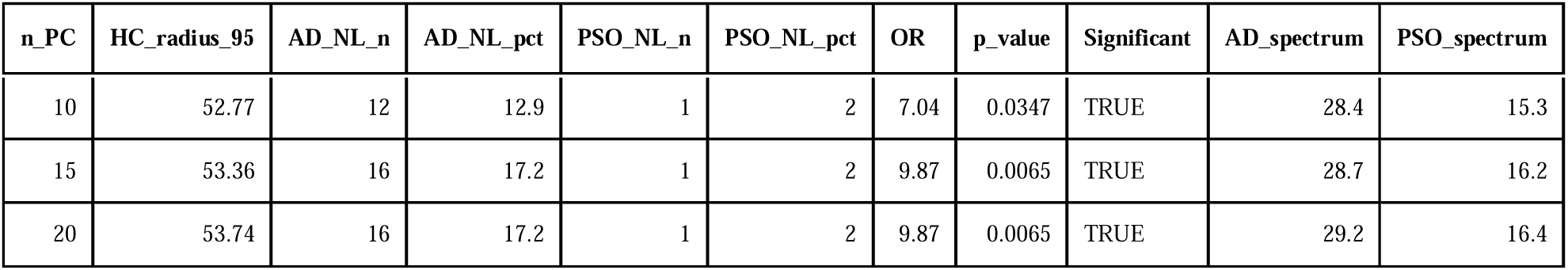
PC dimensionality sensitivity analysis for homeostatic boundary-crossing results at three PC thresholds. At 10 PCs: HC radius = 52.77; AD_NL 12.9% outside (12/93); PSO_NL 2.0% (1/49); OR = 7.04, p = 0.035 (significant). At 15 PCs: HC radius = 53.36; AD_NL 17.2% (16/93); OR = 9.87, p = 0.0065 (significant). At 20 PCs (primary): HC radius = 53.74; AD_NL 17.2% (16/93); OR = 9.87, p = 0.0065 (significant). Spectrum scores: AD_NL 28.4%, 28.7%, 29.2% at 10, 15, 20 PCs respectively; PSO_NL 15.3%, 16.2%, 16.4%. The boundary-crossing finding is statistically significant and directionally consistent across all three PC choices, confirming that the result is not an artefact of the specific dimensionality selected.

**Supplementary Table S3:** Distance method sensitivity analysis comparing Euclidean (primary) and Mahalanobis boundary definitions. Euclidean: AD_NL 17.2% outside, PSO_NL 2.0% outside, OR = 9.87, p = 0.007. Mahalanobis (accounts for covariance structure of healthy reference cloud): AD_NL 52.7% outside, PSO_NL 32.7% outside, OR = 2.28, p = 0.033. Both methods confirm significantly elevated boundary escape in AD_NL versus PSO_NL. The smaller OR under Mahalanobis distance reflects the fact that this metric defines a narrower effective boundary in high-variance PCA dimensions, causing a higher baseline escape rate in all groups; the differential effect (AD_NL > PSO_NL) is replicated. Euclidean distance is used as the primary method because it provides a more interpretable geometric boundary anchored to the observed spread of healthy controls.

| Method | AD_NL_outside_pct | OR | p_value |
| --- | --- | --- | --- |
| **Euclidean (primary)** | 17.2 | 9.87 | 0.007 |
| **Mahalanobis (sensitivity)** | 52.7 | 2.28 | 0.033 |

**Supplementary Table S4:**
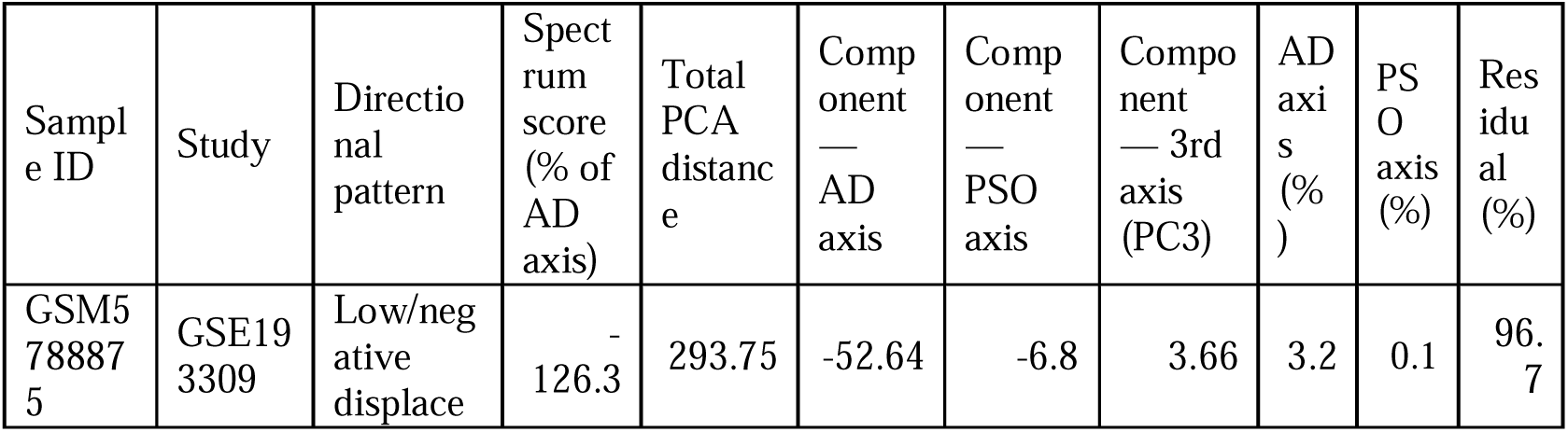
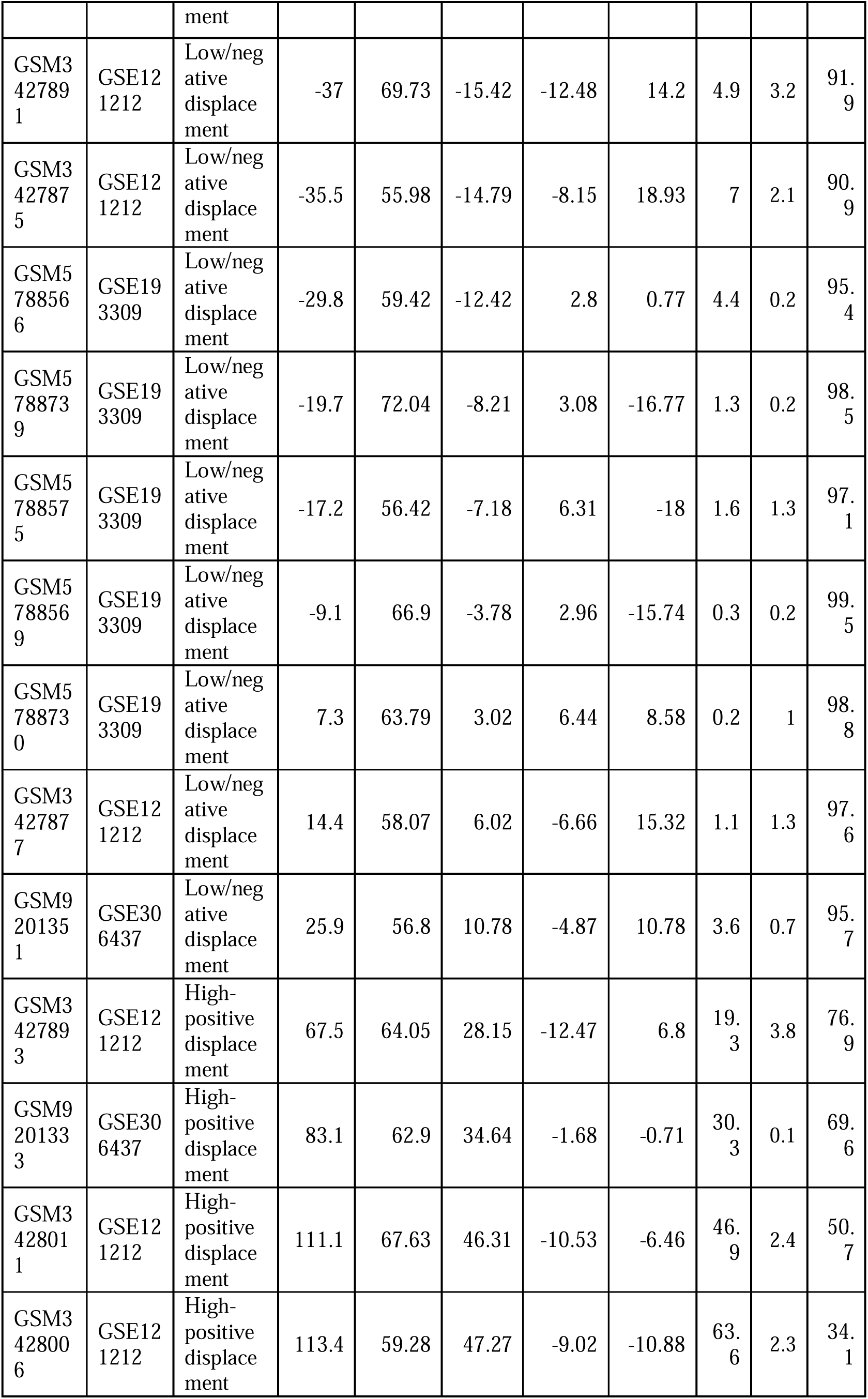
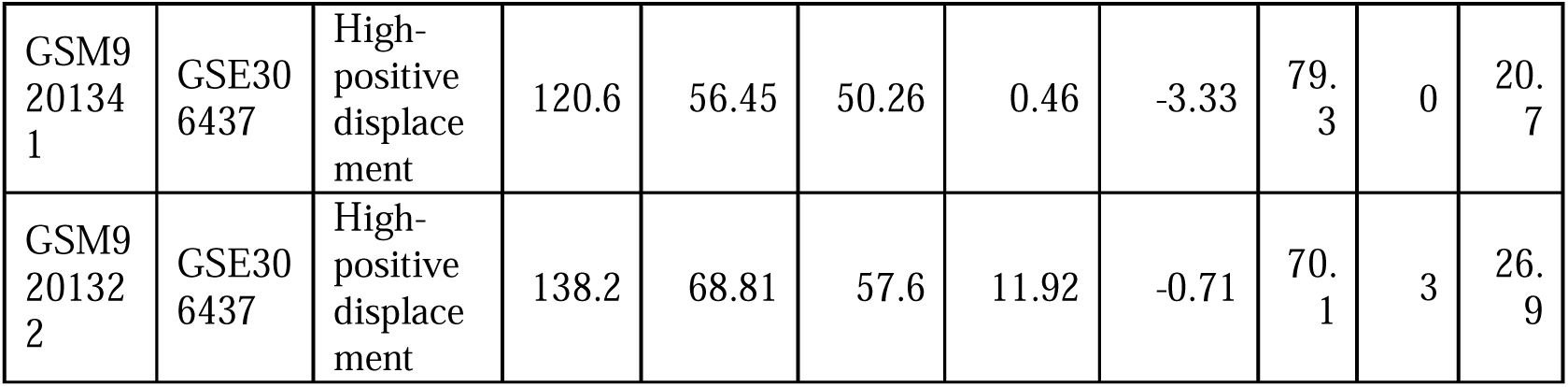
Per-sample axis decomposition for all 16 boundary-crossing AD_NL samples. Columns: sample ID (GSM number), study of origin, directional pattern (low/negative-displacement or high-positive-displacement), spectrum score (% of AD axis distance), total Euclidean PCA distance from healthy centroid, component projections onto the AD axis, PSO axis, and third axis (PC3), and the percentage of total displacement variance attributable to each component (AD axis %, PSO axis %, residual %). Low/negative-displacement samples (n = 10): AD-axis fraction 0.2–4.9%, residual fraction 91.9–99.5%. High-positive-displacement samples (n = 6): AD-axis fraction 19.3–79.3%, residual fraction 20.7–76.9%. The extreme outlier GSM5788875 (spectrum −126.3%, total distance 293.75, residual 96.7%) is included and confirmed not to drive the group-level boundary-crossing result in sensitivity analyses.

